# Meiotic effects of *MSH4* copy number variation support an adaptive role for post-polyploidy gene loss

**DOI:** 10.1101/482521

**Authors:** Adrián Gonzalo, Marie-Odile Lucas, Catherine Marquis, Andrew Lloyd, Eric Jenczewski

**Affiliations:** Institut Jean-Pierre Bourgin, INRA, AgroParisTech, CNRS, Université Paris-Saclay, 78000, Versailles, France; Department of Cell and Developmental Biology, John Innes Centre, Norwich, NR4 7UH, United Kingdom; INRA UMR1349 Institut de Génétique, Environnement et Protection des Plantes, France; Institute of Biological, Environmental, and Rural Sciences, Aberystwyth University, Aberystwyth SY23 3EB, United Kingdom

## Abstract

Many eukaryotes descend from polyploid ancestors that experienced massive duplicate gene loss. This genomic erosion is particularly strong for duplicated (meiotic) recombination genes that return to a single copy more rapidly than genome average following polyploidy. To better understand the evolutionary forces underlying duplicate loss, we analysed how varying copy numbers of *MSH4*, an essential meiotic recombination gene, influences crossover formation in allotetraploid *Brassica napus*. We show that faithful chromosome segregation and crossover frequencies between homologous chromosomes are unchanged with *MSH4* duplicate loss; by contrast, crossovers between homoeologous chromosomes (which result in genomic rearrangements) decrease with reductions in *MSH4* copy number. We also found that inter-homoeologue crossovers originate almost exclusively from the MSH4-dependent crossover pathway. Limiting the efficiency of this pathway by decreasing the copy number of key meiotic recombination genes could therefore contribute to adaptation to polyploidy, by promoting regular chromosome segregation and genomic stability.

## Text

Gene duplication and gene loss are two driving forces in evolution ^1–3^. These two mechanisms are well illustrated by the evolution of the meiotic recombination machinery. At the outset, the emergence of meiosis required key evolutionary breakthroughs ^4–6^ that were all made possible through iterative gene duplications followed by acquisition of new functions. These include, among others, the formation of programmed DNA double-strand breaks by SPO11 proteins ^7^, the promotion of double-strand break repair using homologous templates by DMC1, RAD51 and some other related proteins ^8^ or the resolution of recombination intermediates as crossovers by MSH and MLH proteins ^9^. After this initial phase, genes involved in meiotic recombination (and DNA repair in general) have tended to return preferentially to single copy following subsequent duplications ^10,11^, with some even lost completely in a few specific lineages ^7,12,13^. A major gene duplication pathway particularly prevalent in flowering plants is Whole-Genome Duplication (WGD) ^14^. Although all angiosperms have experienced at least one, and usually multiple rounds of ancient WGDs ^14^, meiotic recombination gene duplicates show a precipitous rate of loss after such duplications, returning to a single copy more rapidly than the genome wide average ^15^.

The importance of gene duplication in powering functional innovation is well understood. In contrast, the adaptive significance of post-WGD duplicate loss, a process referred to as *fractionation* ^16^, is less well supported. Over time, several hypotheses have been proposed, invoking neutral ^15^ or selective ^11,17,18^ processes, but the empirical evidence supporting them is very little. Nevertheless, it is reasonable to assume that the fate of duplicated genes depends, at least in part, on how duplicate loss or persistence affects fitness ^2^. The simplest solution to gain insights into this issue is to evaluate the functional consequences of duplicate gene loss in a recently formed polyploid. Meiotic recombination duplicates are particularly well suited to this approach as meiotic defects cause reduced fertility, a detrimental feature counteracted by natural selection.

Here we describe how meiotic recombination is affected in an allopolyploid (*Brassica napus*) when the functional dosage of a key recombination gene (*MSH4*) decreases. MSH4 is an essential protein for the main (class I) meiotic crossover pathway in most eukaryotes. This pathway, which ensures that at least one obligate crossover is formed between every pair of homologous chromosomes ^19^, is required for faithful chromosome segregation and fertility. *MSH4* is also one of the meiotic recombination genes that return most consistently to single copy after WGD in plants ^15^. *Brassica napus* (AACC, 2n=38) is a recent allotetraploid crop bred for high seed yield that originated from interspecific hybridizations between the ancestors of *B. rapa* (AA, 2n=20) and *B. oleracea* (CC, 2n=18) ∼7,500 years ago ^20^. It is also, and most importantly, one of only two allopolyploid species (along with bread wheat ^21,22^) for which meiosis has been thoroughly analysed ^23^.

Here we show that regular meiosis is achieved in *B. napus* even when the number of functional *MSH4* copies is reduced to a minimum. By contrast, we demonstrate that the number of *MSH4*-dependent crossovers formed between homoeologues (i.e. related chromosomes inherited from different *B. napus* progenitors) fluctuates in a dosage-sensitive way. It is maximum when all copies are functional, decreases progressively with the number of copies and approximates zero when all *MSH4* copies are not functional. This suggests that loss of one *MSH4* copy has no negative impacts and is potentially beneficial for *B. napus* meiosis, which improves when crossovers between homoeologues are suppressed ^23^.

## Results

### *MSH4* illustrates convergent loss of duplicated meiotic recombination genes following WGDs

We first considerably expanded our previous survey (firstly limited to 11 plant genome sequences ^15^) to measure the extent to which *MSH4* duplicates are rapidly lost after WGD. We compiled a taxonomically broad data set spanning 95 fungi, 39 animal and 76 angiosperm species that, together, cover 48 independent WGDs ranging in age from few thousand to almost 500 Million years. As illustrated in Figure 1, we identified only one intact copy of *MSH4* in all species except the very recent polyploids (*Brassica napus*, *Chenopodium quinoa*, *Gossypium hirsutum* and *Zygosaccharomyces rouxii*) that formed <10 000 years ago (in red on figure 1A and 1C) and *Diplocarpon rosae* and *Hortaea werneckii* (in orange on Figure 1A), two fungi that have retained 80% and 95% of gene duplicates post-WGDs, respectively ^24,25^. Remnants of a second copy of *MSH4* carrying inactivating changes were also detected in *Linum usitatissimum*, *Glycine max* and *Populus trichocarpa* (in orange or blue on Figure 1A) as well as in *Salmo salar* and *Oncorhynchus mykiss* (in blue on Figure 1B). *L. usitatissimum* and *G. max* are two of the diploid plant species of our survey that experienced the most recent WGD events: 5-9 and 5-13 million years ago, respectively ^26,27^. *P. trichocarpa* (∼49 MY old WGD ^28^) is a long life cycle perennial species that shows a low rate of duplicate gene loss post-WGD ^15^. The inactivating changes found (important sequence gaps, mutations and truncations in exonic sequences) in those *MSH4* pseudogenes, are likely signatures of ongoing fractionation.

**Figure 1:**
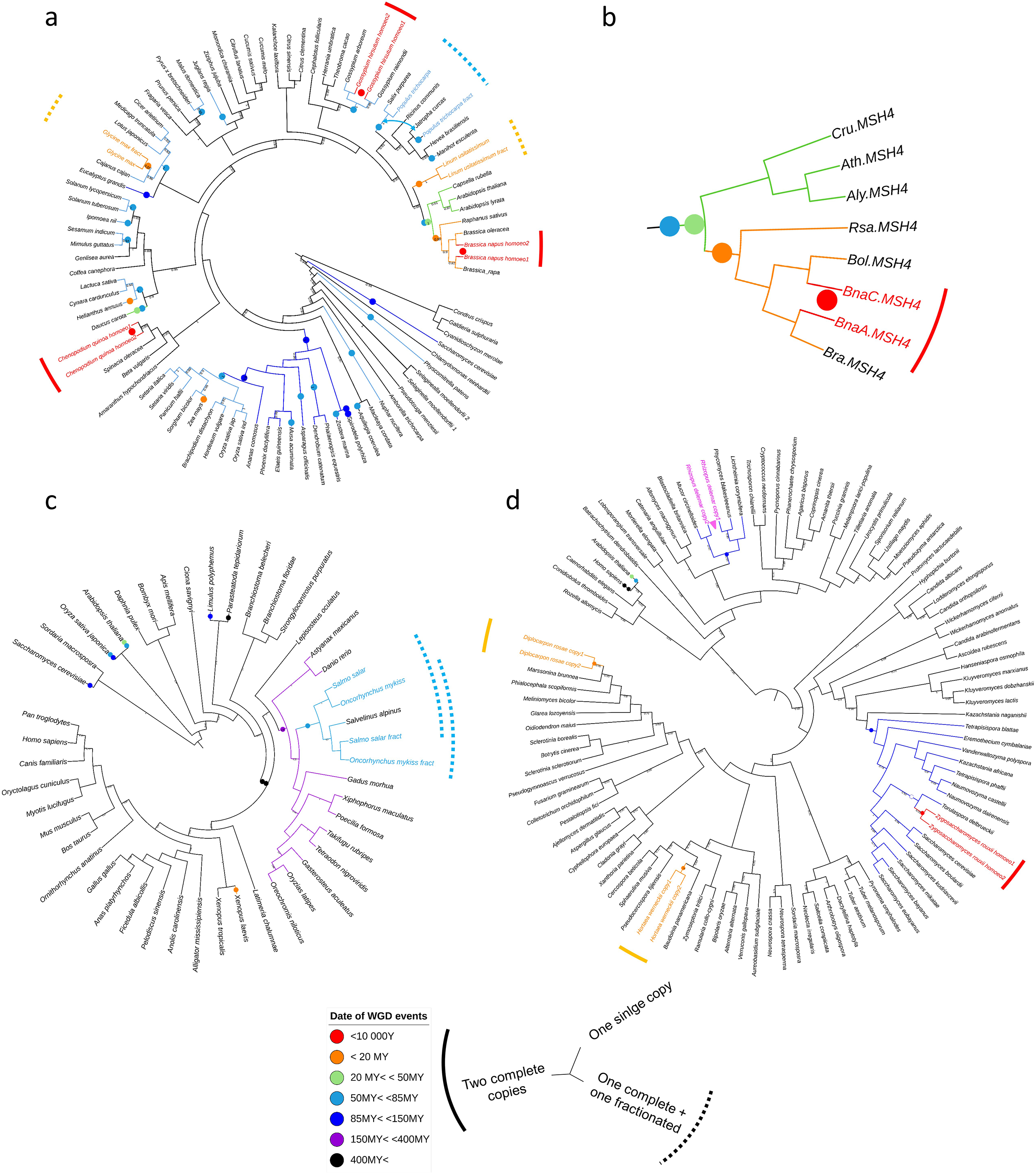
*MSH4* gene copy numbers among angiosperms (a, b), animals (c) and fungi (d) Maximum Likelihood trees based on MSH4 amino acid sequences are provided. For the sake of clarity, species names are indicated instead of gene names (except for b). Note that figure 1b is not simply a zoom in on the previous tree (1a), but a new ML tree based on a new alignment of MSH4 amino acid sequences in the Brassicaceae. In all trees, branch support is given as Shimodaira-Hasegawa-like Likelihood Ratio Test (aLRT SH-like). Coloured disks superimposed along the branches of the trees give the age range for past WGDs. Open disks indicate bursts of gene duplication that have not been formally associated with a WGD event ^29,30^. The pink triangle (and associated *Rhizopus delemar* copies; 1d) represents tandem duplicates. Note that Figure 1c does not reflect all of the WGDs that contributed to the evolution of animals. The reason is that many of the additional unrepresented WGD events were detected using somatic transcriptome data ^31^ from which MSH4 is excluded. Full-length duplicates and recent duplicated with one fractionated copies are written with the color that corresponds to the age of the WGD (i.e. red: <10,000 years; orange: <20MY; light blue: 50MY< < 85MY). Regarding the most recent WGD (i.e. red: <10,000 years), only allopolyploids are represented because there is no way to account for the presence of *MSH4* “duplicates” in extant autopolyploids, which truly correspond to (multiple) real alleles.

All these results confirm that *MSH4* is a textbook case of the rapid loss of meiotic recombination duplicates following WGDs; they also indicate that, although very rapid on an evolutionary timescale (i.e. a few million years), *MSH4*-duplicate loss is not immediate. The presence of *MSH4* duplicates in allopolyploids provides an avenue for evaluating the consequences of *MSH4* duplicate loss.

### *Brassica napus* carries two full-length copies of *MSH4* that are differentially expressed

As shown in Figure 1b, *B. napus* contains two full-length *MSH4* homologues, thereafter referred as to *BnaA.MSH4* and *BnaC.MSH4*. These two copies were inherited from *B. rapa* and *B. oleracea*, respectively, and correspond to homoeologues, *sensu* ^32^. These different genes were identified by querying the *MSH4* coding sequence of *A. thaliana* (AY646927) against the reference assemblies of the three species. A partial (4 exons instead of 24) copy of *MSH4* was also found tandemly arrayed with *BnaA.MSH4* (Supplementary Table 1). This additional copy is not transcribed during meiosis (based on *B. napus* meiotic RNAseq ^33^) and thus not considered for further analyses.

We carried out pyrosequencing to measure the relative mRNA abundance of *BnaA.MSH4* and *BnaC.MSH4* in *B. napus* floral buds. We found that the two copies were transcribed in the three varieties analyzed, with *BnaC.MSH4* contributing most to *MSH4* expression in *B.napus* (Supplementary Table 2); in all varieties, the balance between *BnaA.MSH4* / *BnaC.MSH4* contribution was in the range of 1/6 to 1/3 (Supplementary Table 2). Next, we used *B. napus* meiotic RNAseq data ^33^ to validate the mRNA sequences of *BnaA.MSH4* and *BnaC.MSH4*. The deduced amino acid sequences of *BnaA.MSH4* and *BnaC.MSH4* both contained canonical MutS domains including the highly conserved C-terminal MutSac domain, which is essential for ATPase and DNA-binding activities^34^ (Supplementary Figure 1). This is the region that we targeted to identify EMS-induced mutations affecting MSH4 function.

### TILLING enables production of *msh4* mutants in allotetraploid *B. napus*

Duplicate gene loss may occur via deletion of the (entire or partial) sequence of one duplicate, a process that is still difficult to repeat experimentally, and/or through accumulation of loss-of-function mutations ^3^. In this study, we searched for mutations in *BnaA.MSH4* and *BnaC.MSH4* across 500 M2 plants from the EMS mutagenized RAPTILL population (described in ^35^). Using separate screens for the two genes, we identified 44 and 43-point mutation alleles within 1 kb of the MutSac domain of *BnaA.MSH4* and *BnaC.MSH4*, respectively (Supplementary Figure 2). Amongst these, we selected one mutant allele in *BnaA.MSH4* (hereinafter referred as to *bnaA.msh4-1,* symbolized by A^1^) and two mutant alleles in *BnaC.MSH4* (hereinafter referred as to *bnaC.msh4-1*, symbolized by C^1^, and *bnaC.msh4-2*, symbolized by C^2^, respectively). These alleles were the only ones for which the conserved MutSac domain was disrupted by early stop codons introduced by a point-nonsense mutation (*bnaC.msh4-1*) or because of splice site mutations (*bnaA.msh4-1* and *bnaC.msh4-2*) (Supplementary Figure 2; Supplementary Figure 3).

We then produced two different F1 hybrids that combined *bnaA.msh4-1* with either *bnaC.msh4-1* or *bnaC.msh4-2* (Supplementary Figure 4). These F1s were self-fertilized to produce segregating F2 progenies among which we selected plants that contained varied number and assortments of Wild Type (A^+^ or C^+^) and mutant *msh4* alleles (A^1^, C^1^ or C^2^) (Supplementary Figure 4).

### MSH4 promotes formation of the majority of crossovers between homologues in *B. napus*

In plants as in many other organisms, *msh4* mutants show a severe reduction in crossover leading to the occurrence of numerous univalents (i.e. chromosome that failed to form crossovers) at metaphase I ^36–38^. To ensure that the retained mutations did indeed compromise MSH4 function, we first assessed whether plants carrying only *msh4* mutant alleles (A^1^A^1^C^1^C^1^, A^1^A^1^C^2^C^2^) showed such meiotic defects.

We first analyzed progression of male meiosis in WT *B. napus* cv. *Tanto*, the accession in which *msh4* mutations were identified. Our cytological survey showed that meiosis was very regular in cv. *Tanto*, like in other *B. napus* accessions ^23^. During prophase I, meiotic chromosomes condensed, recombined and underwent synapsis, the close association of two homologous chromosomes via the Synaptonemal Complex (SC), which was complete at pachytene (Supplementary Figure 5a). From diakinesis to metaphase I, 19 discrete bivalents were identifiable in all cells. They consisted of pairs of homologous chromosomes bound together by chiasmata, the cytological manifestation of meiotic crossovers. We estimated that 57% of the bivalents in WT were rings with both arms bound by chiasmata while the remaining 43% were rods with only one arm bound by chiasmata. Assuming that rod and ring bivalents had only one and two crossovers, respectively, we estimated a mean of 29.8 chiasmata per cell in WT *B. napus* cv. *Tanto* (n=27; Table 1). This could be an underestimate however, given that it is not possible to distinguish cytologically single from multiple crossovers clustered on a single arm. The second meiotic division then took place and produced balanced tetrads of four microspores (Supplementary Figure 5f).

In the A^1^A^1^C^1^C^1^ and A^1^A^1^C^2^C^2^ double mutants, the early stages of prophase I were similar to those of WT *B. napus* cv. *Tanto* (Supplementary Figure 5g-h). We confirmed that synapsis proceeded normally in these plants, as demonstrated by immunolocalization of ZYP1 protein, a major component of the central element of the SC ^39^ (Supplementary Figure 6). Meiotic defects became obvious at the end of meiotic prophase when bivalent formation was strongly compromised. We observed a mean number of 20.4-22.6 univalents (70% of chromosomes) that coexisted with 7.7- 8.8 bivalents in both A^1^A^1^C^1^C^1^ and A^1^A^1^C^2^C^2^ (Table 1). Contrary to WT, the majority of bivalents were rods (84%). At metaphase I, the reduction in chiasmata frequency was thus very evident, dropping down to an average of 9.9 chiasmata per cell (Table 1). The univalents then segregated randomly, resulting in unbalanced tetrads (Supplementary Figure 5l).

To test whether the *msh4* mutant alleles completely suppress MSH4 activity, we immuno-localized HEI10 and MLH1, two proteins that specifically mark the sites of MSH4-dependant (i.e. class I) crossovers ^40^. In WT *B. napus* cv. *Tanto*, we consistently counted 29 foci for the HEI10 and MLH1 proteins (Table 1; Figure 2 and 3). By contrast, in the A^1^A^1^C^1^C^1^ and A^1^A^1^C^2^C^2^ double mutants, the residual chiasmata observed were not marked by HEI10 or MLH1 foci (Table 1; Figure 2), suggesting absence of class I crossovers in these plants. This indicated that A^1^A^1^C^1^C^1^ and A^1^A^1^C^2^C^2^ were null *msh4* mutants and that *bnaA.msh4-1*, *bnaC.msh4-1* and *bnaC.msh4-2* alleles encoded nonfunctional MSH4 proteins. We were thus fully equipped to evaluate the consequences of *MSH4* duplicate loss.

**Table 1:** Crossover reduction in absence of functional *MSH4*.

| Genotype | Number of bivalents | Number of chiasmata | Number of MLH1 foci | Number of HEI10 foci |
| --- | --- | --- | --- | --- |
| A<sup>+</sup>A<sup>+</sup>C<sup>+</sup>C<sup>+</sup> | 19 ± 0.01 (n=55) | 29.8 ± 2.02 (n=27) | 29.79 ± 4.78 (n=38) | 29.4 ± 2.62 (n=47) |
| A<sup>1</sup>A<sup>1</sup>C<sup>1</sup>C<sup>1</sup> | 8.8 ± 2.46 (n=89) | 9.89 ± 2.4 (n=89) | 0.49 ± 0.52 (n=19) | 0.6 ± 1.74 (n=39) |
| A<sup>2</sup>A<sup>2</sup>C<sup>2</sup>C<sup>2</sup> | nd | nd | 1.6 ± 1.4 (n=15) | 0.8 ± 1.1 (n=36) |

Data is expressed as mean ± SD. Sample size is given in brackets as the total number of Pollen Mother Cells observed overall biological replicates (when available).

**Figure 2.**
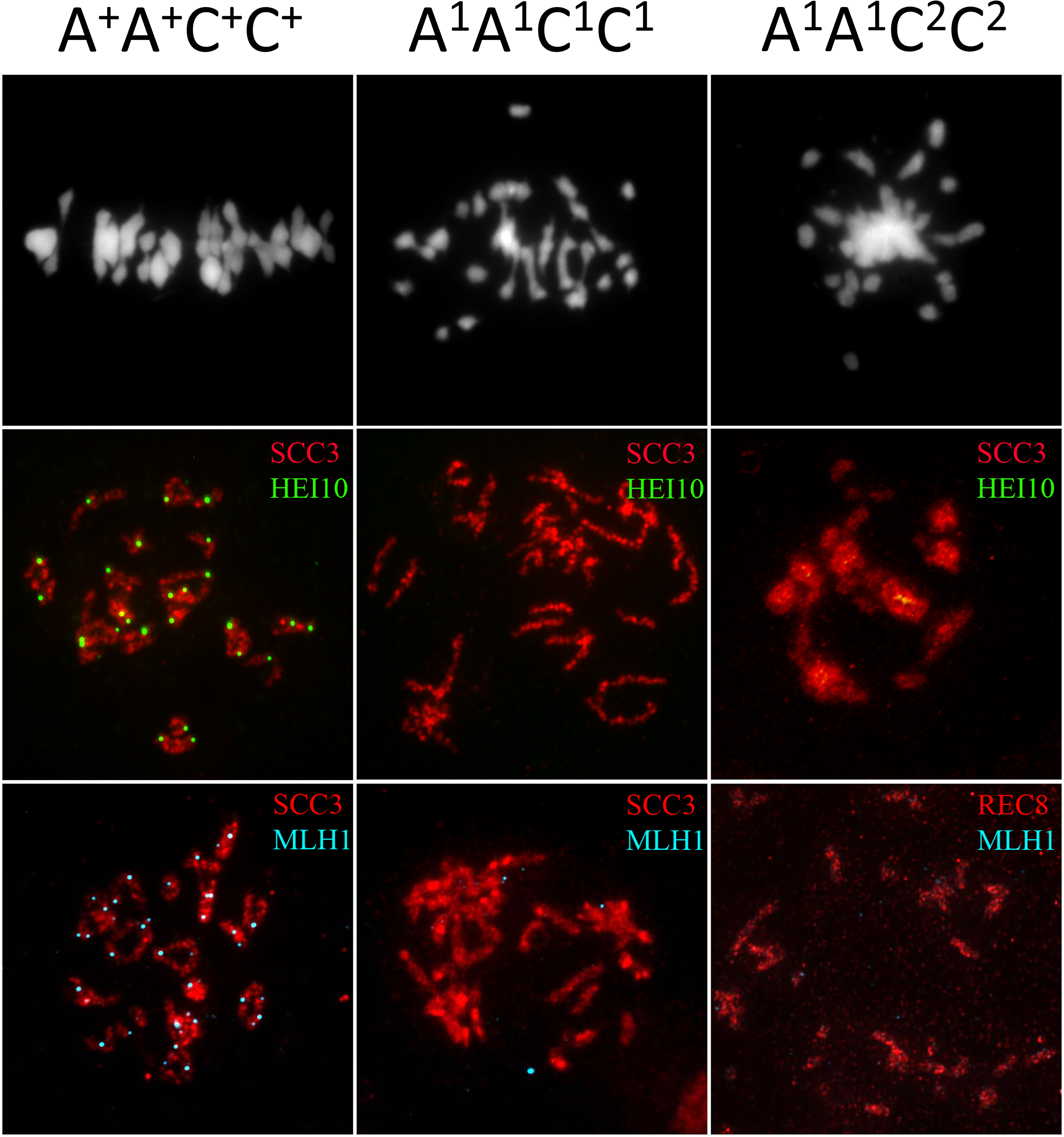
Formation of crossovers between homologous chromosomes in presence and absence of functional *MSH4*. Chiasmata and immunolabeled class I crossovers in WT and two different double mutants combining different *msh4* alleles: A^1^A^1^C^1^C^1^ and A^1^A^1^C^2^C^2^. The upper pictures show DAPI spreads of metaphase I. The middle pictures show dual immunolocalization of SCC3 and HEI10 at diakinesis stage. The lower pictures show dual immunolocalization of SCC3 and MLH1. Bar scale 10 µm.

**Figure 3:**
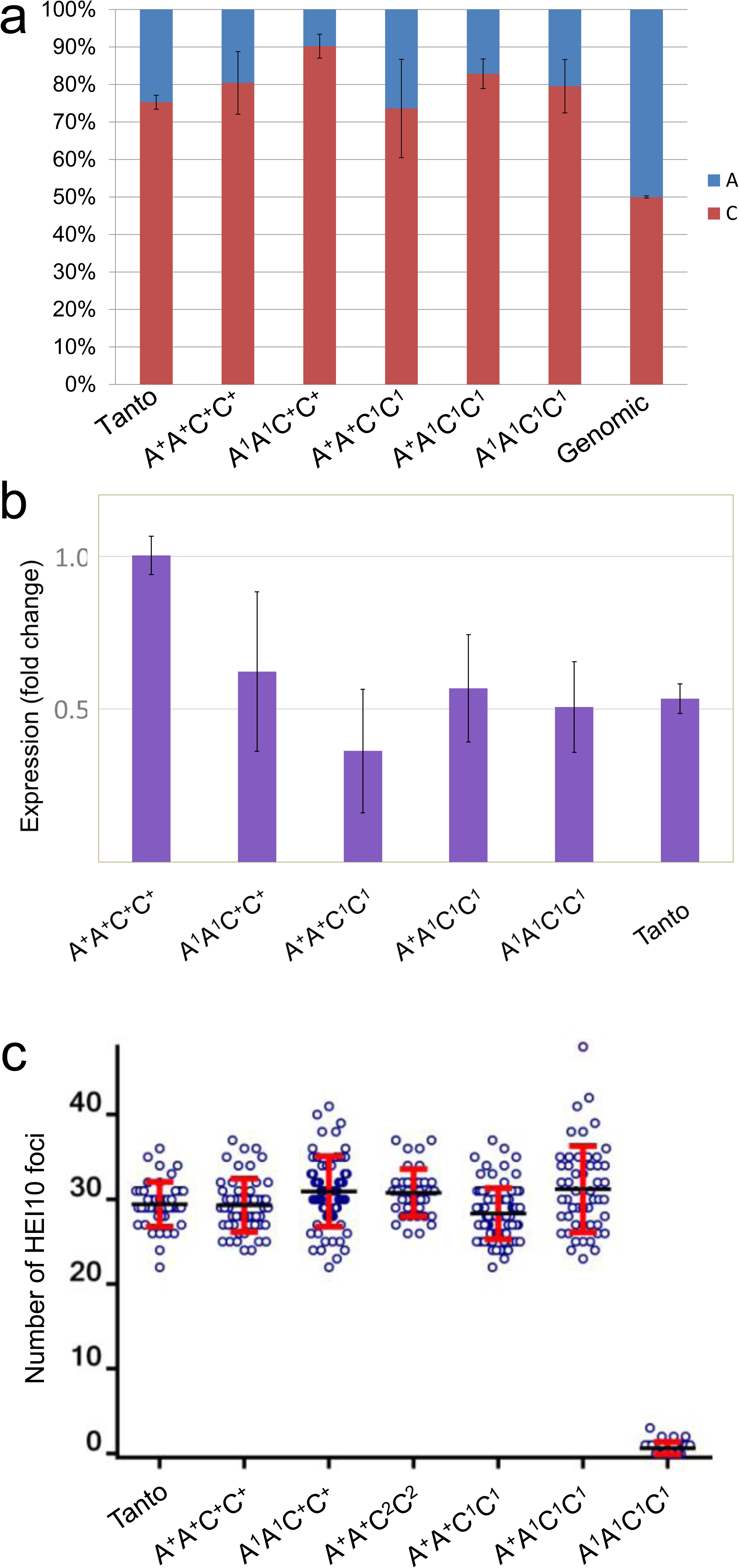
Transcriptional and phenotypic robustness against loss of *MSH4* functional copies. (a) Relative contribution of *BnaA.MSH4* and *BnaC.MSH4* to total MSH4 transcripts evaluated by pyrosequencing. (b) Quantification of the summed expression of *BnaA.MSH4* and *BnaC.MSH4* by real-time PCR, expressed as the normalized fold change (2^-ΔΔCq^) in the target sample relative to the A^+^A^+^C^+^C^+^ genotype. (c) quantification of HEI10 (class I crossover) foci in the different genotypes, as determined by HEI10 immunolocalization.

### *MSH4* copy number does not affect total crossover numbers in *B. napus*

To examine the functional consequences of loss of *MSH4* copies, we characterized meiotic behavior of plants carrying different combinations of WT and mutant alleles (Supplementary Figure 4). This approach assumed that the functional loss of one copy is not compensated by a transcriptional upregulation of the remaining *MSH4* WT alleles. Although quite rare, this phenomenon was not completely unheard of ^41–43^. Owing to the very high similarity between the nucleotide sequences of *BnaA.MSH4* and *BnaC.MSH4*, it was not practically possible to assess the level of transcription of each of the two genes using qPCR alone (whether they show WT or mutant alleles). We thus used pyrosequencing and qPCR to assess the relative contribution and the summed expression, respectively, of *BnaA.MSH4* and *BnaC.MSH4* in the different plants (Figure 3a and 3b; Supplementary Table 2). No significant variation was observed. In all plants showing a combination of WT and mutant alleles, *BnaC.MSH4* remained the most expressed copy while the summed expression of *BnaA.MSH4* and *BnaC.MSH4* was similar to that in WT Tanto. These results suggest that there was neither a strong decay of mutant allele mRNAs, nor an obvious transcriptional compensation between functional and mutant MSH4 alleles.

We then moved on to the cytological survey of the exact same plants. We observed that functional loss of *BnaA.MSH4*, the least expressed *MSH4* copy, or *BnaC.MSH4*, the most expressed *MSH4* copy (Supplementary Table 2, figure 3c), all resulted in a WT-like meiosis (Figure 3c; Supplementary Figure 7 and Supplementary Figure 8). The A^1^A^1^C^+^C^+^, A^+^A^+^C^1^C^1^ and A^+^A^+^C^2^C^2^ plants all showed 19 bivalents at metaphase I (Supplementary Figure 7) and the same number of HEI10 foci as the WT (Figure 3c; Supplementary Figure 8). These results indicated that, irrespective of their unequal transcriptional contribution, *BnaA.MSH4* and *BnaC.MSH4* are both equally functional and able to complement one another. We then explored the extreme situation where *BnaC.MSH4* was completely depleted while only one allele of the *BnaA.MSH4* was functional. This plant, i.e. A^+^A^1^C^1^C^1^, carried the minimum functional *MSH4* dosage possible, with only one functional allele of the least expressed copy (e.g. > 90 % reduction in functional transcript). Despite this minimal composition, A^+^A^1^C^1^C^1^ showed exclusive bivalent configuration and about 31 HEI10 foci per meiocyte as in WT (Figure 3c; Supplementary Figure 7f and Supplementary Figure 8f).

All these results indicated that normal class I crossover formation is not sensitive to *MSH4* duplicate loss, providing that (at least) one functional copy of *MSH4* is present in the plant. There should therefore be no obvious selective force opposing *MSH4* duplicate loss in *B. napus*.

### Crossover formation between homoeologues is strongly affected by MSH4 copy number

Although some crossovers can occasionally form between homoeologous chromosomes in *B. napus* ^33,44^, their frequency remains very low ^45–47^, making comparison of their occurrence between the different genetic backgrounds very difficult. To overcome this limitation, and test whether crossover formation between homoeologues is robust against loss of functional *MSH4* copies, we produced allohaploid plants (AC) through microspore culture. These plants contain one unique copy of each of the 19 *B. napus* chromosomes (n=19) and thus no longer have homologous chromosomes. Despite the absence of homologue pairs, some crossovers are formed between homoeologous partners. To study how *msh4* mutations affect these crossover events, we used two different F1 hybrids (A^+^A^1^C^+^C^1^ and A^+^A^1^C^+^C^2^) to produce a series of allohaploid plants with different combinations of WT and mutant alleles (Supplementary Figure 4). As in euploids, we first used pyrosequencing and qPCR to verify that there was no transcriptional compensation between functional and mutant *MSH4* alleles. As in euploids, we did not observe significant changes in the summed expression of A- and C-copies of *MSH4*, nor in their relative contributions, between plants showing varied number and assortments of WT (A^+^ or C^+^) and mutant msh4 alleles (A^1^, C^1^ or C^2^) (Figure 4ab).

**Figure 4:**
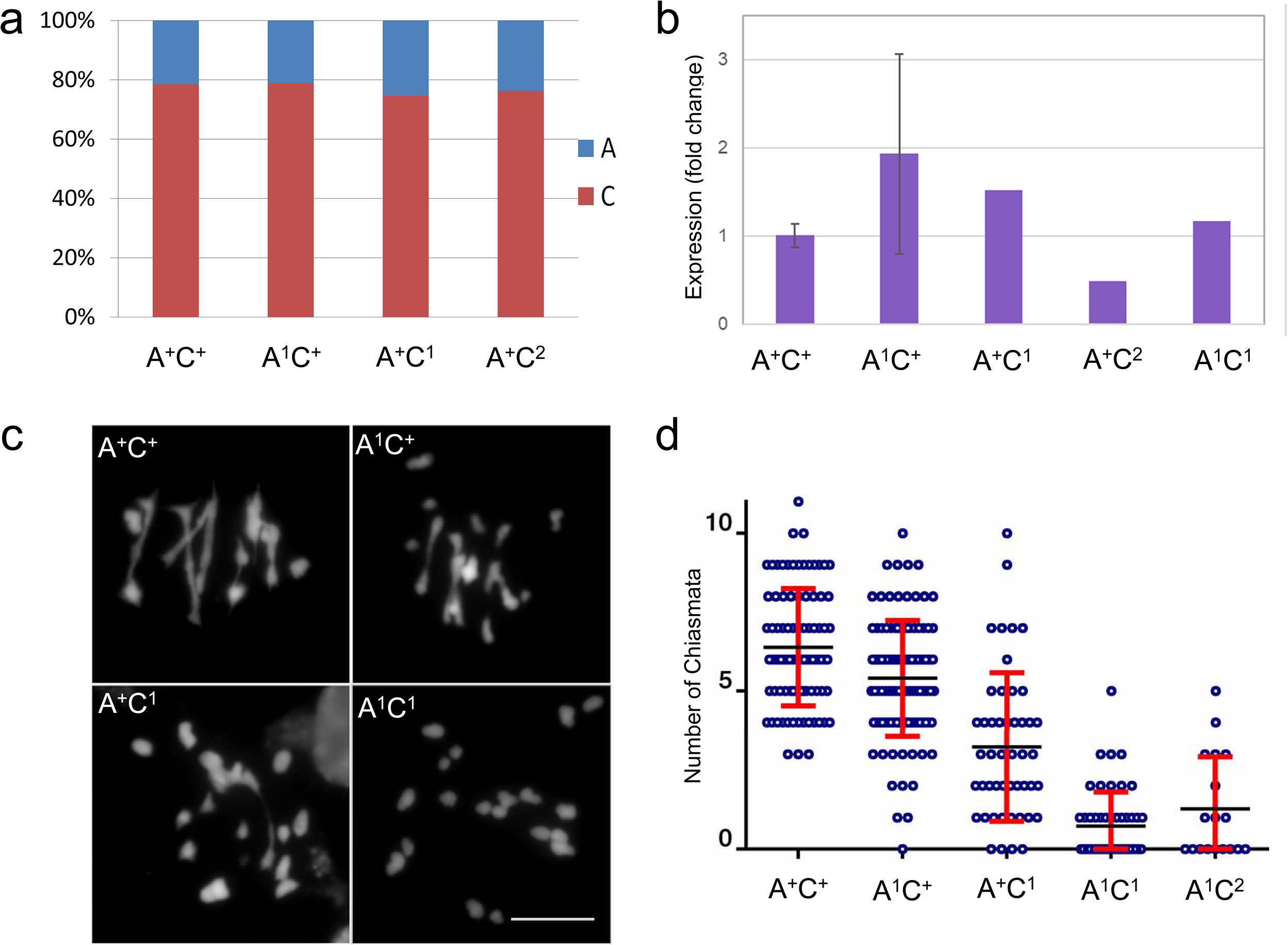
Transcriptional and phenotypic consequences of *MSH4* duplicate loss among *Brassica napus* allohaploids. (a) Relative contribution of *BnaA.MSH4* and *BnaC.MSH4* to total MSH4 transcripts evaluated by pyrosequencing. (b) Quantification of the summed expression of *BnaA.MSH4* and *BnaC.MSH4* by real-time PCR, expressed as the normalized fold change (2^-ΔΔCq^) in the target sample relative to the A^+^C^+^ genotype. (c) Chromosome associations at Metaphase I and (d) quantification of HEI10 (class I crossover) foci in the different genotypes.

We then counted chiasmata to estimate and compare crossover frequencies between the different plants. WT allohaploids (A^+^C^+^) showed on average 6.4 chiasmata distributed over a mean number of 1 ring and 4.4 rod bivalents (Figure 4c and 4d; Supplementary Table 3). Interestingly, we observed a slight but significant decrease in chiasmata frequency (compared to WT allohaploids t=3.7, p-value= 0.0003) when the least expressed copy was depleted; the mean number of chiasmata in A^1^C^+^ plants dropped down to 5.4 crossovers (Figure 4c and 4d; Supplementary Table 3). A stronger and significant reduction in chiasmata was observed when the most expressed copy was depleted, down to 2.16 chiasmata per cell in A^+^C^1^ (compared to A^1^C^+^; t=9.35, p-value < 0.0001) and 2.67 chiasmata per cell in A^+^C^2^ (compared to A^1^C^+^; t=4.51, p-value= 0.0001) allohaploids, respectively (Figure 4c and 4d; Supplementary Table 3). In these two plants, almost all bivalents were rods and only very rare ring bivalents were observed. Finally, when both *MSH4* copies were depleted, the number of residual chiasmata was further decreased, down to 0.73 in A^2^C^1^ (compared to A^+^C1; t=4.51, p-value < 0.0001) and 0.9 in A^2^C^2^ (Figure 4c and 4d; Supplementary Table 3). Contrary to the euploids, immunolocalization of MLH1 and HEI10 proteins could not be used to confirm the decay of class I crossovers in all these plants. We previously showed that in *B. napus* allohaploid meiosis the two proteins do not always co-localize during meiosis in *B. napus* allohaploid ^23^, which made Class I crossover estimates technically impossible. This notwithstanding, our results indicated that inter- homoeologue class I crossover formation is sensitive to *MSH4* duplicate loss (Figure 4d).

## Discussion

The pervasiveness of gene loss during evolution ^2,48^ could be explained either because it may drive adaptive innovations ^2,3^ or simply because functionally redundant, duplicated (and dosage-insensitive) genes are dispensable ^49–51^. To evaluate each of these two possibilities, we interrogated the (rapid) return of *MSH4* duplicates to singletons following whole genome duplication events ^15^.MSH4 accounts for the majority of meiotic crossovers in most organisms, including plants ^36,52^. We first confirmed that *MSH4* plays the same role in recent allotetraploid *B. napus* as in other plants: though not required for synapsis (Supplementary Figure 6), MSH4 is essential to ensure normal crossover numbers between homologues (Table 1; Figure 2 and 3) and, therefore, strictly required to ensure fertility.

Most importantly, our results showed that normal levels of homologous crossovers are robust against *MSH4* gene duplicate loss. Irrespective of the copy-specific expression level, functional loss of one copy or the other did not result in any reduction of class I crossovers between homologues. This holds true even when only one single functional allele of the least expressed copy was present (in the A^+^A^1^C^1^C^1^plant, Figure 4; Supplementary Figure 7 and Supplementary 8). This dispensability of *MSH4* duplicates (as long as one functional allele is present), could lead the hypothesis that this gene returns to a single copy stochastically, as degenerative mutations randomly affect either one of the two copies. We did observe, however, that *MSH4* genes returned to singletons in all species, except the most recent polyploids (Figure 1). *MSH4* duplicate loss was effective within a few million years irrespective of whether the species are ancient autopolyploids (e.g. soybean, poplar) or ancient allopolyploids (e.g. maize). According to some authors ^17,53^, such a consistent pattern of gene loss is unlikely to be the result of a purely random and neutral process, but rather meet the expectation for deterministic duplicate loss ^10,17,54,55^. Our results on inter- homoeologue crossovers in allohaploid plants showing different combinations of WT and mutant alleles may support this second scenario.

It has long been acknowledged that homoeologous recombination disrupts faithful chromosome segregation and must be kept to a minimum to optimize fertility in allopolyploids ^56,57^. Providing inter-homoeologue crossover formation in allohaploids is a good proxy of what is happening in the euploids, where both homologues and homoeologues are competing, our results show that those illegitimate events are strongly dependent on MSH4 and that they can be sharply reduced by decreasing *MSH4* dosage. As describe above, this reduction can occur without altering crossover formation between homologues, another prerequisite for faithful chromosome segregation. The reasons for this intriguing specificity of dosage sensitivity are unknown. It could be tentatively linked to the abortion of a high number of late inter-homoeologue recombination intermediates that are formed downstream the MSH4-dependent pathway but do not eventually mature into crossovers. This phenomenon has been observed in *B. napus* allohaploids ^21,23^ and in wheat ^21^; it may actually only represent the final step in a broader abortion process, which required a greater cellular concentration of MSH4 to stabilize inherently less stable / more transient early inter-homoeologue recombination intermediates. Future work should test this mechanistic hypothesis. This notwithstanding, since loss of one *MSH4* copy does not affect the homologous crossovers, its specific effect on the illegitimate events could only improve regular chromosome segregation (hence optimizing fertility) and could thus be subject to positive selection. This advantageous return of *MSH4* to singleton would illustrate the “Less-is-more Hypothesis” ^2^ in its polyploid variant: the “Selected Single Copy Hypothesis” ^10,53^ which postulates that some maladaptive effects of polyploidy might be buffered by decreasing the dosage of the genes involved. This also echoes recent findings in yeast interspecific hybrids, which regained fertility thanks to duplicate gene loss ^58,59^.

On a broader scale, our results shed light on the longstanding conundrum of meiotic adaptation in allopolyploids. They indicate that reducing the efficiency of the main recombination (class I) pathway could be sufficient to bias the choice of recombination partners away from the homoeologues, thus promoting inter-homologue crossovers. This idea is in line with recent results from wheat, in which an extra copy of *ZIP4*, a gene involved in the class I crossover pathway, was recently shown to be instrumental in preventing inter-homoeologue crossovers in wheat ^60^. The mode of action of this extra copy is not yet known but it is likely to modulate the class I pathway ^21^. Counter to the situation in wheat, we here show that alteration of class I crossovers can also occur via gene loss, due to degenerative mutational events that knock-out one copy of a pair of homoeologues. Thus, this research could open new directions to better appreciate the evolutionary role of gene loss ^2,3^ in the context of polyploidy, where this phenomenon is pervasive ^48^.

## Methods

### Plant material

*Brassica napus* L. cv. *Tanto* is a double-low spring cultivar obtained at INRA Rennes (France) that was used to develop the RAPTILL EMS population ^35^ in which we looked for mutations in *BnaA.MSH4* and *BnaC.MSH4*. Microspore culture was performed to isolate allohaploid plants following the protocol described in ^61^. All plants were cultivated in standard long day greenhouse conditions (16-hour light/8-hour night photoperiod, at 22°C day and 18°C night).

### TILLING

We screened the RAPTILL EMS population following exactly the same protocol as described in ^35^. Briefly, we targeted a region of 1 Kb of the MutSac domain of *BnaA.MSH4* and *BnaC.MSH4.* We used copy-specific primers (T_MSH4AF1, T_MSH4AR1, T_MSH4CF1 and T_MSH4CR1; Supplementary Table 4) to amplify *BnaA.MSH4* and *BnaC.MSH4* separately. The resulting amplicons span the regions between the positions 4144-5126 and 4161-5148 in the genomic sequences of *BnaA.MSH4* and *BnaC.MSH4*, respectively. The screens for mutations then implemented the PMM (Plant Mutated on its Metabolites) method ^62,63^.

### Genotyping

Molecular screening for *MSH4* alleles involved direct Sanger sequencing of PCR product (using T_MSH4AF1 and T_MSH4AR1 primers; Supplementary Table 4) for *BnaA.msh4-1* allele by. The other mutant alleles were identified using Cleaved Amplified Polymorphic Sequences (CAPS) assay targeting the causative EMS-SNP. The list of primers and restriction enzymes in given in Supplementary Table 4 and Supplementary Table 5.

### Cytological analysis

Staged anthers were isolated from fixed buds in ethanol:acetic acid 3:1 (v/v). They were used to produce meiotic spreads according to ^64^. Chiasma number was estimated on metaphase I spread chromosomes counterstained with 4’,6-diamidino-2-phenylindole (DAPI) based on bivalent shape. Open “rod” bivalents were considered to contain one single chiasma while closed ring bivalents were scored as two (one on each arm). For the study of allohaploids, all countings were done blindly. Spreads were also used for immunolocalization.

### Immunolocalization of meiotic proteins

Immunodetection of MLH1, HEI10, REC8 and SCC3 proteins was performed following the protocol described by ^65^. Immunodetection of ASY1 and ZYP1 followed the protocol described by^66^ that uses fresh anthers. The anti-MLH1 and anti-HEI10 antibodies, both obtained in rabbit serum, were used at dilution 1:200. The antibodies raised against SCC3 and REC8, from guinea pig (GP) and rat serum, respectively, were diluted at 1:250. Anti-ASY1 and anti-ZYP1, obtained in guinea pig (GP) and rabbit, respectively, were both used at dilution 1:500. Secondary antibodies were Alexa488- anti-rabit (green) and Alexa568-anti-GP and –anti-rat (red).

### Fluorescent microscopy

All images were obtained using a Zeiss AxioObserver microscope and were analyzed by means of Zeiss Zen software and were organized in panels with Adobe Photosoft CS3.

### RNA extraction and cDNA obtention

RNA was extracted from buds selected by size using RNeasy Mini Kit (Qiagen). One extra step of clean-up and DNAse treatment was added to the extraction protocol. Resulting RNA was used for cDNA obtention using RevertAid First Strand cDNA Synthesis Kit (Thermo Fisher Scientific). The presence and absence of cDNA and residual genomic DNA was assessed using the primers Q_UBC21R1, Q_UBC21L1, Q_MSH4F1 and Q_MSH4R1 (Supplementary Table 4). Euploid and allohaploid plants were sampled simultaneously under same conditions.

### Quantitative PCR

Three biological replicates were analyzed for every euploid genotype, each replicate being located on a separate qPCR plate. Intercalibration between plates was performed using two internal controls common to all three plates. The nine allohaploid plants analyzed were also distributed among three different plates, but the same actual number of biological replicates could not be achieved for all plants. Three technical replicates were used for all (euploid and allohaploid) plants, all located onto the same plate. We used the ubiquitin gene *UBC21* as a reference to normalize the expression of the target *MSH4*. The primers used in these experiments (Q_UBC21R1, Q_UBC21L1, Q_MSH4F1 and Q_MSH4R1) are given in Supplementary Table 4.

### Pyrosequencing

Pyrosequencing reactions were designed with primers flanking homeologue specific SNPs in the coding sequence of *BnaA.MSH4* and *BnaC.MSH4*, as described in ^15^. Pyrosequencing was performed on meiotic cDNA, and on genomic DNA as a control to normalize the ratios against possible pyrosequencing biases. Pyro sequencing reaction was performed with PyroMark Q24 v2.0.6 (Qiagen).

### Statistical analysis

The relative expression of a *MSH4* genes (compared to *UBC21*) was calculated based on ‘delta delta Ct’ (ΔΔCt) values. One-way ANOVA was then performed using R to test for mutant genotype differences. ANOVA tests to analyze cytological data were performed using Prism GraphPad.

### Phylogeny construction

Gene trees were reconstructed using the Maximum-likelihood (ML) method implemented in the phylogeny pipeline provided by Phylogeny.fr (http://www.phylogeny.fr/). Multiple Alignment were carried out with MUSCLE (full mode). Alignment curation was implemented using GBLOCKS, allowing smaller final blocks and gap positions within the final blocks. Phylogenetic trees were generated using PhyML v3.0 ^67^ after estimating the Gamma distribution parameters, the proportion of invariables sites and the Transition/transversion ratio. The sequences of the MSH4 amino acid sequences are provided in Supplementary Table 6. The trees were customized and annotated using iTOL (https://itol.embl.de/). The occurrence and age for the past WGDs were taken from ^24,25,30,68–73^.

## Supporting information

## Acknowledgments

We would like to thank Karine Alix, Christine Mézard, Raphaël Mercier and Mathilde Grelon for critical reading and discussion of the manuscript. We also thank Fatiha Benyahya for her help and acknowledge AELRED for performing the TILLING experiment in *B. napus*. This work was funded through the ANR project ANR-14-CE19-0004 – CROC. The IJPB benefits from the support of the LabEx Saclay Plant Sciences-SPS (ANR-10-LABX-0040-SPS). A.G. was funded by the Marie-Curie “COMREC” network FP7 ITN-606956. A.L. was funded by the International Outgoing Fellowships PIOF-GA-2013-628128 POLYMEIO.

## Author contributions

Study design: A.G., A.L. and E.J.; Plant material production: A.G., M.O.L., C.M.; Cytological analyses: A.G.; transcriptome analyses: A.G., A.L.; Phylogenetic and statistical analyses: E.J.; A.G., A.L. and E.J. contributed to writing the manuscript.

## Materials & Correspondence

Eric Jenczewski is the author to whom correspondence and material requests should be addressed.

## References

1. Ohno, S. Evolution by Gene Duplication. (New York Springer) (1970).

2. Albalat, R. & Cañestro, C. Evolution by gene loss. Nat. Rev. Genet. 17, 379–391 (2016).

3. Sharma, V. et al. A genomics approach reveals insights into the importance of gene losses for mammalian adaptations. Nat. Commun. 9, 1–9 (2018).

4. Villeneuve, A. & Hillers, K. J. Whence meiosis? Cell 106, 647–650 (2001).

5. Marcon, E. & Moens, P. B. The evolution of meiosis: recruitment and modification of somatic DNA-repair proteins. BioEssays 27, 795–808 (2005).

6. Wilkins, A. S. & Holliday, R. The evolution of meiosis from mitosis. Genetics 181, 3–12 (2009).

7. Malik, S.-B., Ramesh, M. A., Hulstrand, A. M. & Logsdon, J. M. Protist homologs of the meiotic Spo11 gene and topoisomerase VI reveal an evolutionary history of gene duplication and lineage-specific loss. Mol. Biol. Evol. 24, 2827–41 (2007).

8. Lin, Z., Kong, H., Nei, M. & Ma, H. Origins and evolution of the recA/RAD51 gene family: evidence for ancient gene duplication and endosymbiotic gene transfer. Proc. Natl. Acad. Sci. U. S. A. 103, 10328–10333 (2006).

9. Lin, Z., Nei, M. & Ma, H. The origins and early evolution of DNA mismatch repair genes--multiple horizontal gene transfers and co-evolution. Nucleic Acids Res. 35, 7591–603 (2007).

10. Paterson, A. H. et al. Many gene and domain families have convergent fates following independent whole-genome duplication events in Arabidopsis, Oryza, Saccharomyces and Tetraodon. Trends Genet. 22, (2006).

11. De Smet, R. et al. Convergent gene loss following gene and genome duplications creates single-copy families in flowering plants. Proc. Natl. Acad. Sci. 110, 2898–2903 (2013).

12. Kohl, K. P., Jones, C. D. & Sekelsky, J. Evolution of an MCM complex in flies that promotes meiotic crossovers by blocking BLM helicase. Science 338, 1363–5 (2012).

13. Fernandes, J. B. et al. FIGL1 and its novel partner FLIP form a conserved complex that regulates homologous recombination. PLoS Genet. 14, e1007317 (2018).

14. Jiao, Y. et al. Ancestral polyploidy in seed plants and angiosperms. (2011). DOI:10.1038/nature09916

15. Lloyd, A. H. et al. Meiotic gene evolution: Can you teach a new dog new tricks? Mol. Biol. Evol. 31, 1724–1727 (2014).

16. Langham, R. J. et al. Genomic duplication, fractionation and the origin of regulatory novelty. Genetics 166, 935–45 (2004).

17. Li, Z. et al. Gene Duplicability of Core Genes Is Highly Consistent across All Angiosperms. Plant Cell 28, 326–44 (2016).

18. Emery, M. et al. Preferential retention of genes from one parental genome after polyploidy illustrates the nature and scope of the genomic conflicts induced by hybridization. PLoS Genet. 14, e1007267 (2018).

19. Krishnaprasad, G. N. et al. Variation in crossover frequencies perturb crossover assurance without affecting meiotic chromosome segregation in Saccharomyces cerevisiae. Genetics 199, 399–412 (2015).

20. Chalhoub, B. et al. Early allopolyploid evolution in the post-Neolithic Brassica napus oilseed genome. Science (80-.). 345, 950–953 (2014).

21. Martín, A. C., Shaw, P., Phillips, D., Reader, S. & Moore, G. Licensing MLH1 sites for crossover during meiosis. Nat. Commun. 5, 4580 (2014).

22. Martín, A. C., Rey, M.-D., Shaw, P. & Moore, G. Dual effect of the wheat Ph1 locus on chromosome synapsis and crossover. Chromosoma (2017). DOI:10.1007/s00412-017-0630-0

23. Grandont, L. et al. Homoeologous Chromosome Sorting and Progression of Meiotic Recombination in Brassica napus: Ploidy Does Matter! Plant Cell 26, 1448–1463 (2014).

24. Neu, E., Featherston, J., Rees, J. & Debener, T. A draft genome sequence of the rose black spot fungus Diplocarpon rosae reveals a high degree of genome duplication. PLoS One 12, e0185310 (2017).

25. Sinha, S. et al. Insight into the Recent Genome Duplication of the Halophilic Yeast Hortaea werneckii: Combining an Improved Genome with Gene Expression and Chromatin Structure. G3 (Bethesda). 7, 2015–2022 (2017).

26. Schmutz, J. et al. Genome sequence of the palaeopolyploid soybean. Nature 463, 178–83 (2010).

27. Wang, Z. et al. The genome of flax (Linum usitatissimum) assembled de novo from short shotgun sequence reads. Plant J. 72, 461–73 (2012).

28. Tuskan, G. A. et al. The genome of black cottonwood, Populus trichocarpa (Torr. & Gray). Science 313, 1596–604 (2006).

29. van Hooff, J. J. E., Snel, B. & Seidl, M. F. Small homologous blocks in phytophthora genomes do not point to an ancient whole-genome duplication. Genome Biol. Evol. 6, 1079–85 (2014).

30. Marcet-Houben, M. & Gabaldón, T. Beyond the Whole-Genome Duplication: Phylogenetic Evidence for an Ancient Interspecies Hybridization in the Baker’s Yeast Lineage. PLoS Biol. 13, e1002220 (2015).

31. Li, Z. et al. Multiple large-scale gene and genome duplications during the evolution of hexapods. Proc. Natl. Acad. Sci. U. S. A. 115, 4713–4718 (2018).

32. Glover, N. M., Redestig, H. & Dessimoz, C. Homoeologs : What Are They and How Do We Infer Them? Trends Plant Sci. 21, 609–621 (2016).

33. Lloyd, A. et al. Homoeologous exchanges cause extensive dosage-dependent gene expression changes in an allopolyploid crop. New Phytol. 217, 367–377 (2018).

34. Tachiki, H. et al. Domain organization and functional analysis of Thermus thermophilus MutS protein. Nucleic Acids Res. 26, 4153–9 (1998).

35. Blary, A. et al. FANCM Limits Meiotic Crossovers in Brassica Crops. Front. Plant Sci. 9, 1–13 (2018).

36. Higgins, J. D., Armstrong, S. J., Franklin, F. C. H. & Jones, G. H. The Arabidopsis MutS homolog AtMSH4 functions at an early step in recombination: Evidence for two classes of recombination in Arabidopsis. Genes Dev. 18, 2557–2570 (2004).

37. Zhang, L. et al. Crossover formation during rice meiosis relies on interaction of OsMSH4 and OsMSH5. Genetics 1–35 (2014). DOI:10.1534/genetics.114.168732

38. Wang, C. et al. The role of OsMSH4 in male and female gamete development in rice meiosis. J. Exp. Bot. 67, 1447–1459 (2016).

39. Higgins, J. D., Sanchez-moran, E., Armstrong, S. J., Jones, G. H. & Franklin, F. C. H. The Arabidopsis synaptonemal complex protein ZYP1 is required for chromosome synapsis and normal fidelity of crossing over. 2488–2500 (2005). DOI:10.1101/gad.354705.ments

40. Chelysheva, L. A., Grandont, L. & Grelon, M. Immunolocalization of meiotic proteins in Brassicaceae: method 1. Methods Mol. Biol. 990, 93–101 (2013).

41. Wong, S. L. & Roth, F. P. Transcriptional compensation for gene loss plays a minor role in maintaining genetic robustness in Saccharomyces cerevisiae. Genetics 171, 829–33 (2005).

42. Springer, M., Weissman, J. S. & Kirschner, M. W. A general lack of compensation for gene dosage in yeast. Mol. Syst. Biol. 6, 368 (2010).

43. DeLuna, A., Springer, M., Kirschner, M. W. & Kishony, R. Need-based up-regulation of protein levels in response to deletion of their duplicate genes. PLoS Biol. 8, e1000347 (2010).

44. Higgins, E. E., Clarke, W. E., Howell, E. C., Armstrong, S. J. & Parkin, I. A. P. Detecting de Novo Homoeologous Recombination Events in Cultivated Brassica napus Using a Genome-Wide SNP Array. G3 (Bethesda). (2018). DOI:10.1534/g3.118.200118

45. Sharpe, A. G., Parkin, I. A. P., Keith, D. J. & Lydiate, D. J. Frequent nonreciprocal translocations in the amphidiploid genome of oilseed rape (Brassica napus). Genome 38, 1112–1121 (1995).

46. Udall, J., Quijada, P. & Osborn, T. Detection of chromosomal rearrangements derived from homologous recombination in four mapping populations of Brassica napus L. Genetics 169, 967–979 (2005).

47. Howell, E. C., Kearsey, M. J., Jones, G. H., King, G. J. & Armstrong, S. J. A and C Genome Distinction and Chromosome Identification in Brassica napus by Sequential Fluorescence in Situ Hybridization and Genomic in Situ Hybridization. Genetics 1857, 1849–1857 (2008).

48. Cheng, F. et al. Gene retention, fractionation and subgenome differences in polyploid plants. Nat. plants 4, 258–268 (2018).

49. Nowak, M. A., Boerlijst, M. C., Cooke, J. & Smith, J. M. Evolution of genetic redundancy. Nature 388, 167–170 (1997).

50. Lynch, M. & Conery, J. The evolutionary demography of duplicate genes. J Struct Funct Genomics 3(1-4), 35–44 (2003).

51. Conant, G. C., Birchler, J. A. & Pires, J. C. Dosage, duplication, and diploidization: clarifying the interplay of multiple models for duplicate gene evolution over time. Curr. Opin. Plant Biol. 19, 91–8 (2014).

52. Mercier, R., Mézard, C., Jenczewski, E., Macaisne, N. & Grelon, M. The Molecular Biology of Meiosis in Plants. Annu. Rev. Plant Biol. 66, 297–327 (2015).

53. Edger, P. P. & Pires, J. C. Gene and genome duplications : the impact of dosage-sensitivity on the fate of nuclear genes. Chromosom. Res 699–717 (2009). DOI:10.1007/s10577-009-9055-9

54. Duarte, J. M. et al. Identification of shared single copy nuclear genes in *Arabidopsis, Populus, Vitis* and *Oryza* and their phylogenetic utility across various taxonomic levels. BMC Evol. Biol. 10, 61 (2010).

55. De Smet, R. et al. Convergent gene loss following gene and genome duplications creates single-copy families in flowering plants. Proc. Natl. Acad. Sci. 110, 2898–2903 (2013).

56. Szadkowski, E. et al. The first meiosis of resynthesized Brassica napus, a genome blender. 102–112 (2010).

57. Gaeta, R. T. & Pires, J. C. Homoeologous recombination in allopolyploids : the polyploid ratchet. New Phytol 18–28 (2010).

58. Ortiz-Merino, R. A. et al. Evolutionary restoration of fertility in an interspecies hybrid yeast, by whole-genome duplication after a failed mating-type switch. PLoS Biol. 15, e2002128 (2017).

59. Braun-Galleani, S., Ortiz-Merino, R. A., Wu, Q., Xu, Y. & Wolfe, K. H. Zygosaccharomyces pseudobailii, another yeast interspecies hybrid that regained fertility by damaging one of its MAT loci. FEMS Yeast Res. 18, (2018).

60. Rey, M.-D. et al. Exploiting The ZIP4 Homologue Within The Wheat Ph1 Locus Has Identified Two Lines Exhibiting Homoeologous Crossover In Wheat-Wild Relative Hybrids. Mol. Breed. 37–95, (2017).

61. Jenczewski, E. et al. PrBn, a major gene controlling homeologous pairing in oilseed rape (Brassica napus) haploids. Genetics 164, 645–653 (2003).

62. Triques, K. et al. Characterization of Arabidopsis thaliana mismatch specific endonucleases: Application to mutation discovery by TILLING in pea. Plant J. 51, 1116–1125 (2007).

63. Triques, K. et al. Mutation detection using ENDO1: Application to disease diagnostics in humans and TILLING and Eco-TILLING in plants. BMC Mol. Biol. 9, 42 (2008).

64. Chelysheva, L. et al. An easy protocol for studying chromatin and recombination protein dynamics during Arabidopsisthaliana meiosis: Immunodetection of cohesins, histones and MLH1. Cytogenet. Genome Res. 129, 143–153 (2010).

65. Chelysheva, L., Grandont, L. & Grelon, M. Immunolocalization of meiotic proteins in Brassicaceae: method 1. Methods Mol Biol 990:93 (2013).

66. Chelysheva, L. et al. Zip4/Spo22 is required for class I CO formation but not for synapsis completion in Arabidopsis thaliana. PLoS Genet. 3, 802–813 (2007).

67. Guindon, S., Delsuc, F., Dufayard, J.-F. & Gascuel, O. Estimating maximum likelihood phylogenies with PhyML. Methods Mol. Biol. 537, 113–37 (2009).

68. Van De Peer, Y., Mizrachi, E. & Marchal, K. The evolutionary significance of polyploidy. Nat Rev. (2017). DOI:10.1038/nrg.2017.26

69. Alix, K., Gérard, P. R., Schwarzacher, T. & Heslop-Harrison, J. S. P. Polyploidy and interspecific hybridization: partners for adaptation, speciation and evolution in plants. Ann. Bot. 120, 183–194 (2017).

70. Berthelot, C. et al. The rainbow trout genome provides novel insights into evolution after whole-genome duplication in vertebrates. Nat. Commun. 5, 3657 (2014).

71. Session, A. M. et al. Genome evolution in the allotetraploid frog Xenopus laevis. Nature 538, 336–343 (2016).

72. Corrochano, L. M. et al. Expansion of Signal Transduction Pathways in Fungi by Extensive Genome Duplication. Curr. Biol. 26, 1577–1584 (2016).

73. Schwager, E. E. et al. The house spider genome reveals an ancient whole-genome duplication during arachnid evolution. BMC Biol. 15, 62 (2017).

