## Supplementary material for "Meiotic effects of *MSH4* copy number variation support an adaptive role for post-polyploidy gene loss"

### Supplementary figure legends

**Supplementary Figure 1. MSH4 proteins in Brassica.** Amino acid sequence alignment of *BnaA.MSH4* and *BnaC.MSH4* with their *A. thaliana* ortholog (*AtMSH4*).

**Supplementary Figure 2: Overview of the *msh4* mutations identified by TILLING in *Brassica napus*.** (a) Schematic representation of *BnaA.MSH4* and *BnaC.MSH4* exon-intron structure identifying the MutSd (green) and MutSac (red) domains (as predicted using the SMART online blast tool SMART). A zoom on the region of the MutSac domain where mutations were searched for is provided for each of the two genes. An arrowhead marks the position of all identified mutations, including *bnaA.msh4-1* (light red), *bnaC.msh4-1* (dark red) and *BnaC.msh4-2* (light red). (b) Distribution of mutations in *BnaA.MSH4* and *BnaC.MSH4*, respectively, according to their type.

**Supplementary Figure 3: Splicing variation in *bnaA.msh4-1* and *bnaC.msh4-2*.** (a) shows a schematic representation of the position of primers used to assess the splicing pattern associated with *bnaA.msh4-1* and *bnaC.msh4-2* mutations, respectively. Black arrows represent the consensus primers that we used to test for the presence of genomic DNA (gDNA). Blue and red arrows represent the pairs of A- and C-copy specific primers, respectively. Amplicon specificity was verified by Sanger sequencing (not shown here). (b) shows the resulting PCR amplicons obtained using genomic DNA (gDNA), cDNA obtained from meiotic buds in Tanto (WT cDNA) and the double mutant $A^1A^1C^2C^2$ (mut cDNA). Blank cDNA (no template for RT) and water are used as negative controls for PCR reactions. One band was produced by *bnaA.msh4-1*, which is very similar in size than that of the wt. Sanger sequencing revealed that this band actually contains two amplicons that show 1 and 16 short deletions compared to the wt sequence, respectively (not shown). By contrast, three bands were obtained using C-copy specific primers and wt cDNA. Only two of these fragments were amplified using mut cDNA, the missing band (arrowhead) corresponding to the correctly spliced mRNA (arrow). (c) shows the alignment of the predicted amino acid sequences encoded by the two aberrant splice variants produced by *bnaA.msh4-1* along with the WT AA sequence. The two spliced variants lead to frameshifts, each followed by premature stop codons.

**Supplementary Figure 4: Genealogy of the plants used to evaluate the consequences of *MSH4* duplicate loss.** Plants homozygous for *bnaA.msh4-1*, *bnaC.msh4-1* and *bnaC.msh4-2* were selected within the corresponding M2 families (from the RapTill population) and crossed to produce two different F1 hybrids. These F1s were self-fertilized to produce F2 progenies among which, plants containing varied number and assortments of Wild Type (A+ or C+) and mutant *msh4* alleles (A1, C1 or C2) were selected. The red lines indicate isolation of allohaploid plants through microspore culture.

**Supplementary Figure 5: Progression of meiosis in *Brassica napus* cv. *Tanto* and *msh4* double mutants.** Comparative DAPI staining of wild type (**a** to **f;** *B.napus* cv *Tanto*) and $A^1A^1C^1C^1$ *msh4* double mutant (**g** to **m**). Different meiotic stages are illustrated: pachytene (**a** and **g**), diplotene (**b** and **h**), diakinesis (**c** and **i**), metaphase I (**d** and **j**), Anaphase/telophase I (**e** and **k**), Late anaphase II (**f** and **l**). Bar scale 10 µm.

**Supplementary Figure 6: Normal Synaptonemal Complex formation during meiosis in *MSH4*-deficient $A^1A^1C^1C^1$ *B.napus*.** Immunolocalization of AtASY1 (green; axial/lateral elements) and AtZYP1 (red; central element) antibodies to spread *B.napus* Pollen Mother Cells. (a) late leptotene showing the axial element of every chromosome marked by ASY1. (b) zygotene showing incipient synapsis marked by the first ZYP1 tracts. (c) late/mid zygotene showing the progression of synapsis, which is eventually achieved at pachytene when the ZYP1 signal is continuous (d).

**Supplementary Figure 7: Meiotic regularity in *Brassica napus* plants combining wild type ($A^+$, $C^+$) and mutant ($A^1$, $C^1$, $C^2$) alleles of *MSH4*.** DAPI spreads of metaphase I obtained from Tanto (**a**), $A^+A^+C^+C^+$ (**b**), $A^1A^2C^+C^+$(**c**), $A^+A^+C^1C^1$ (**d**), $A^+A^+C^2C^2$ (**e**), $A^+A^2C^1C^1$ (**f**), $A^2A^2C^1C^1$ (**g**), $A^1A^1C^2C^2$(**h**). Bar scale 10 µm.

**Supplementary Figure 8: HEI10-dependent crossovers in *Brassica napus* plants combining wild type ($A^+$, $C^+$) and mutant ($A^1$, $C^1$, $C^2$) alleles of *MSH4*.** HEI10 immunolocalization pictures in Tanto (**a**), $A^+A^+C^+C^+$ (**b**), $A^1A^1C^+C^+$(**c**), $A^+A^+C^2C^2$ (**d**), $A^+A^+C^1C^1$ (**e**), $A^+A^1C^1C^1$(**f**), $A^1A^1C^1C^1$(**g**) and $A^2A^2C^1C^1$ (**h**). Bar scale 10 µm.

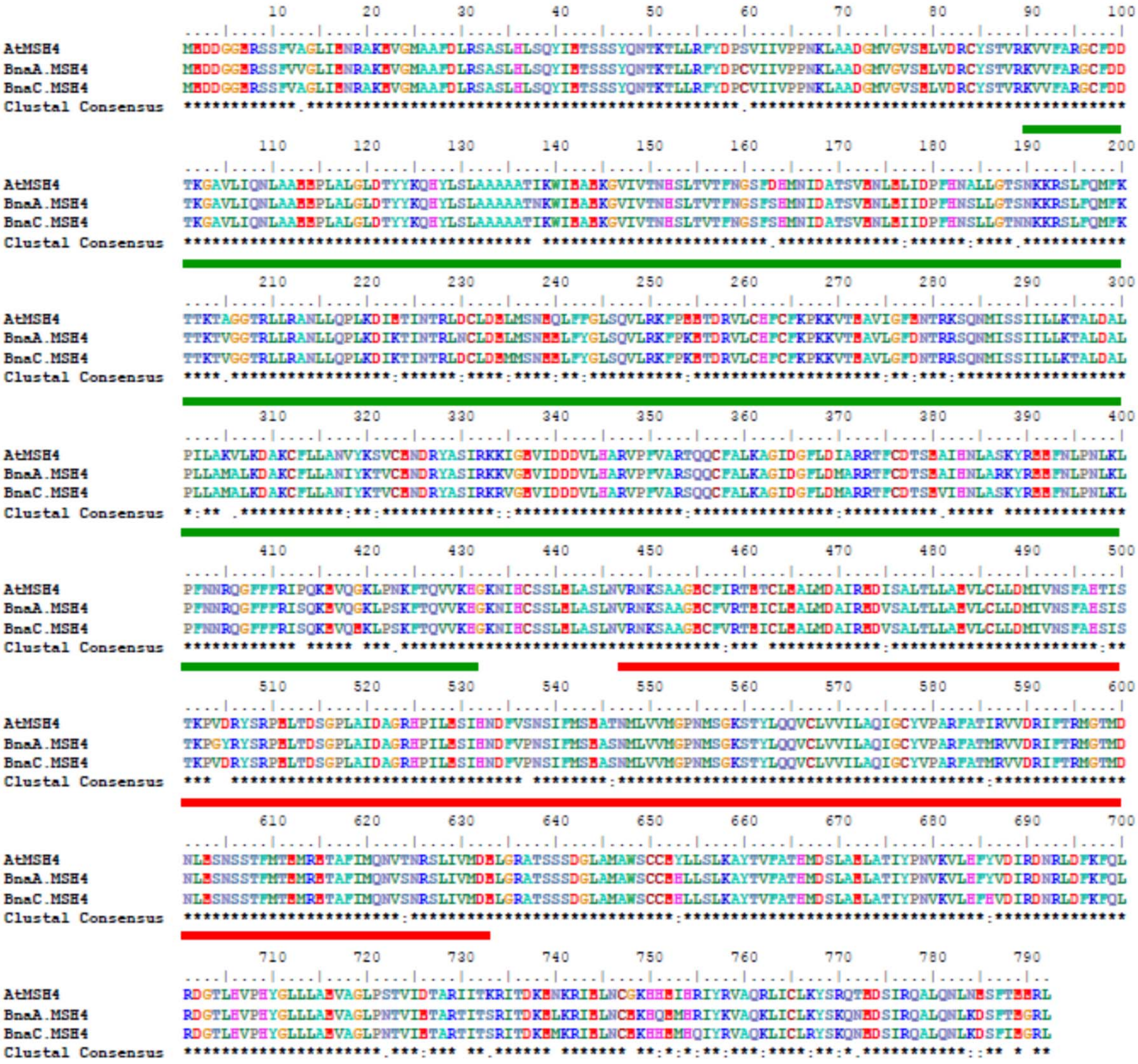


Supplementary Figure 1

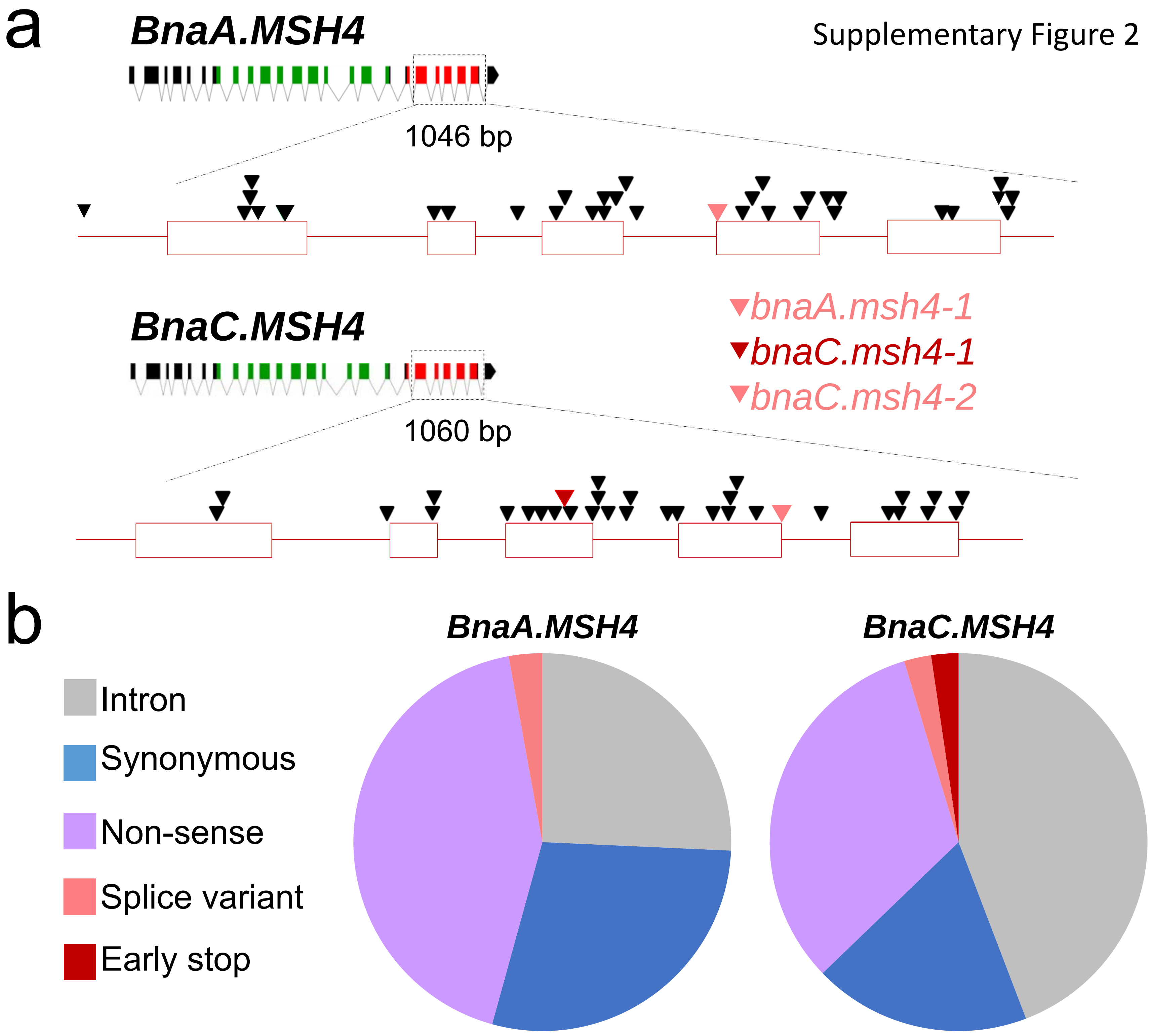
a
BnaA.MSH4
Supplementary Figure 2
1046 bp
bnaA.msh4-1
bnaC.msh4-1
bnaC.msh4-2
BnaC.MSH4
1060 bp
b
BnaA.MSH4
BnaC.MSH4
Intron
Synonymous
Non-sense
Splice variant
Early stop

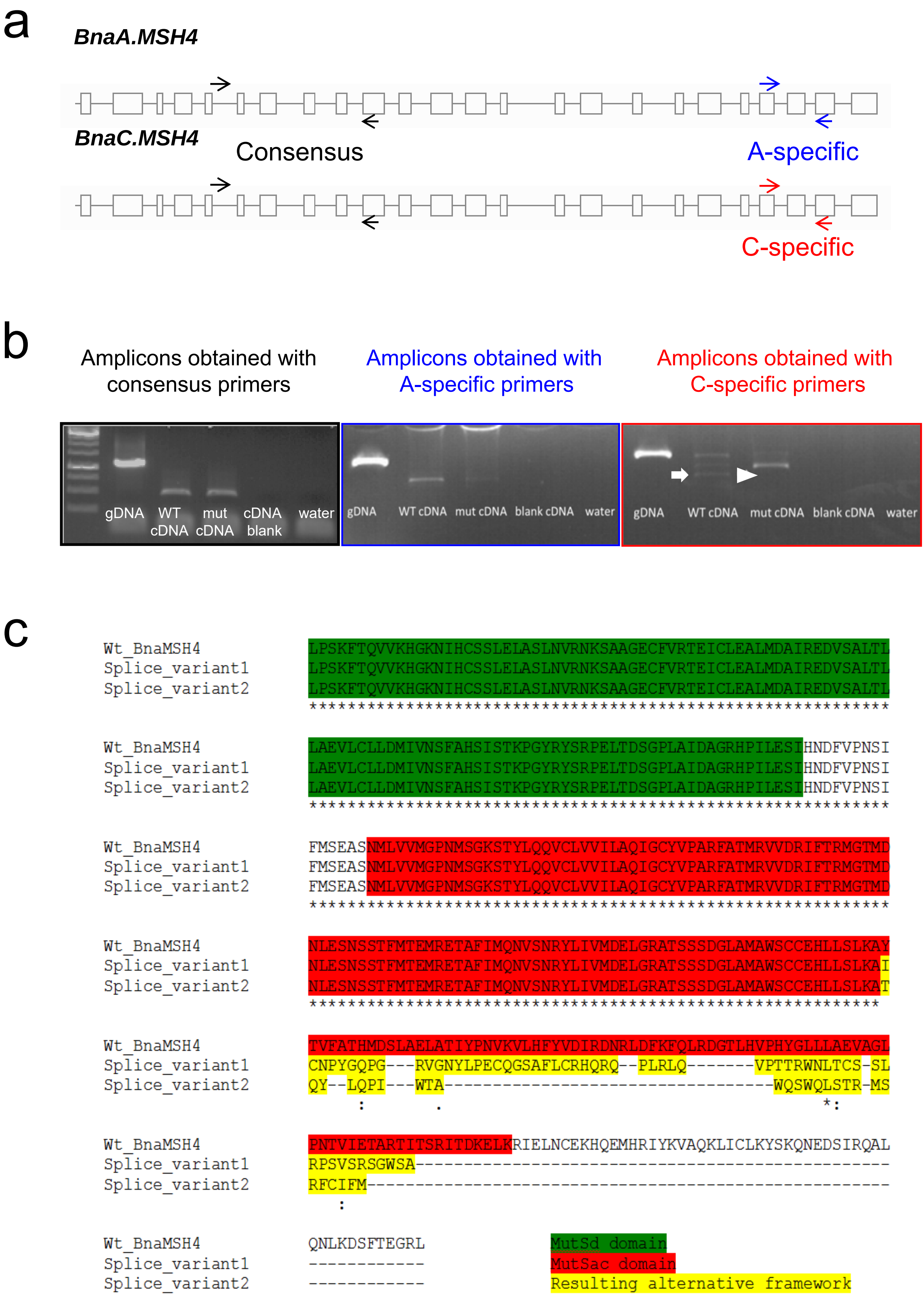


Supplementary Figure 3

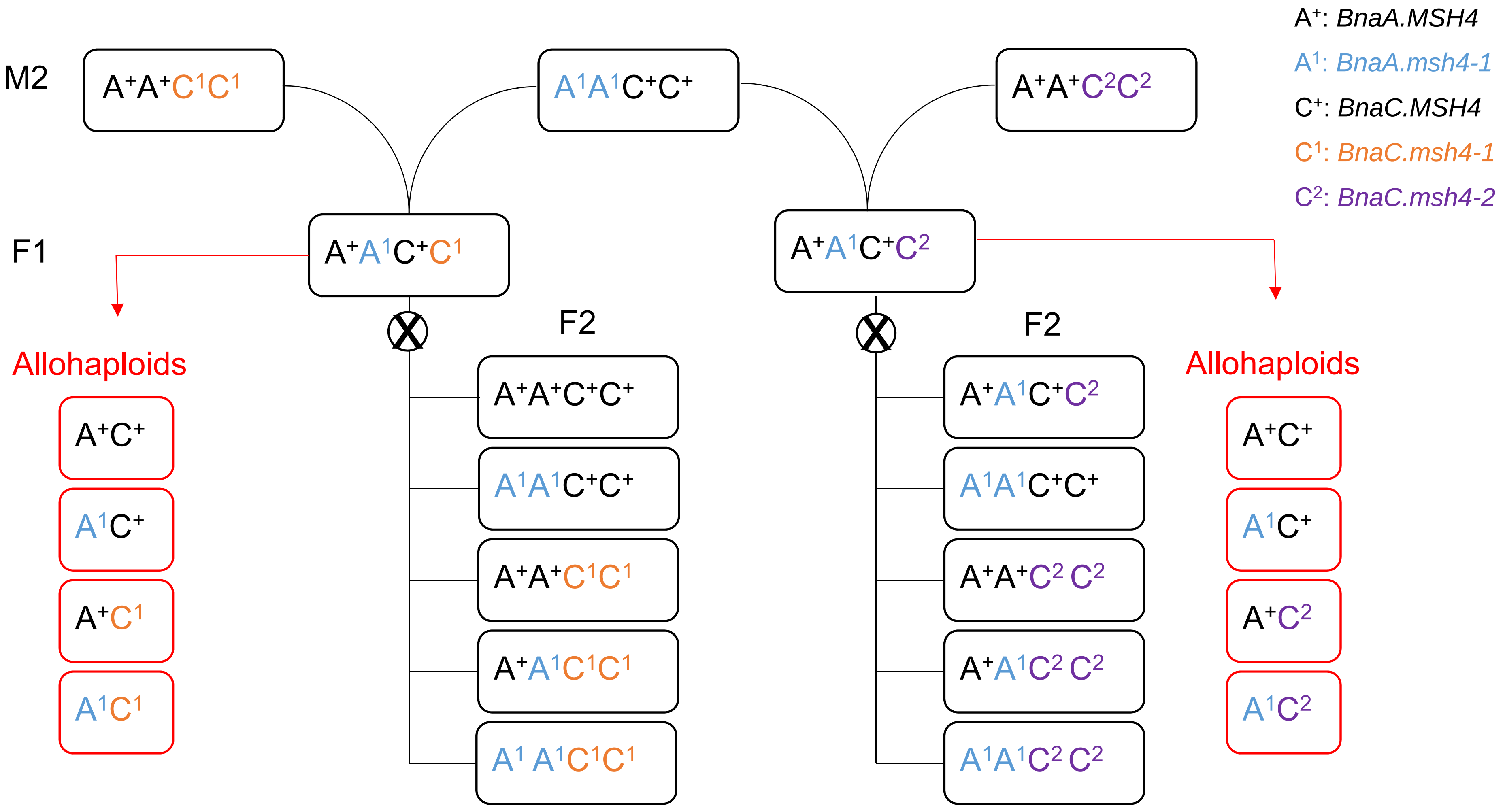


Supplementary Figure 4

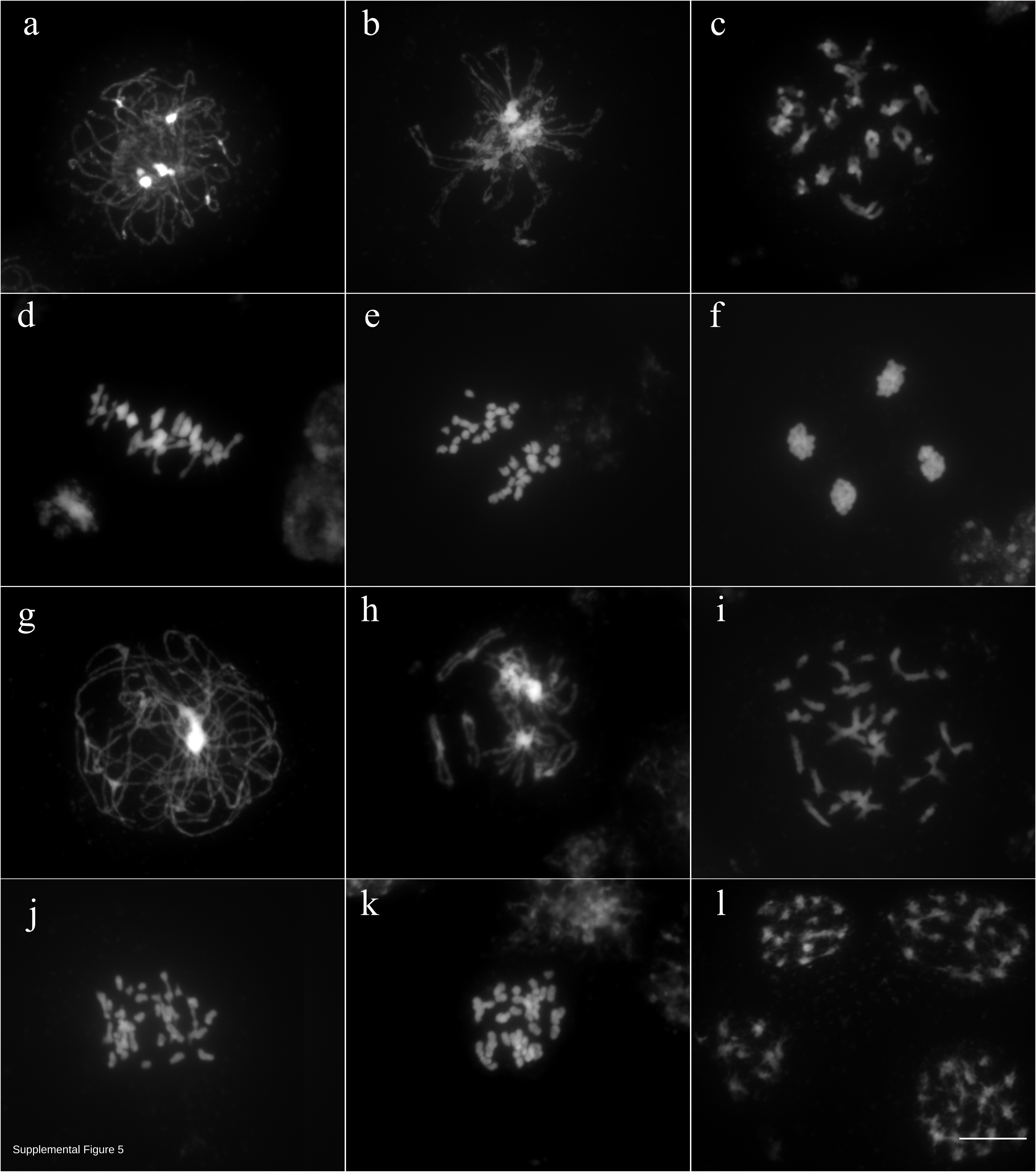

Supplemental Figure 5

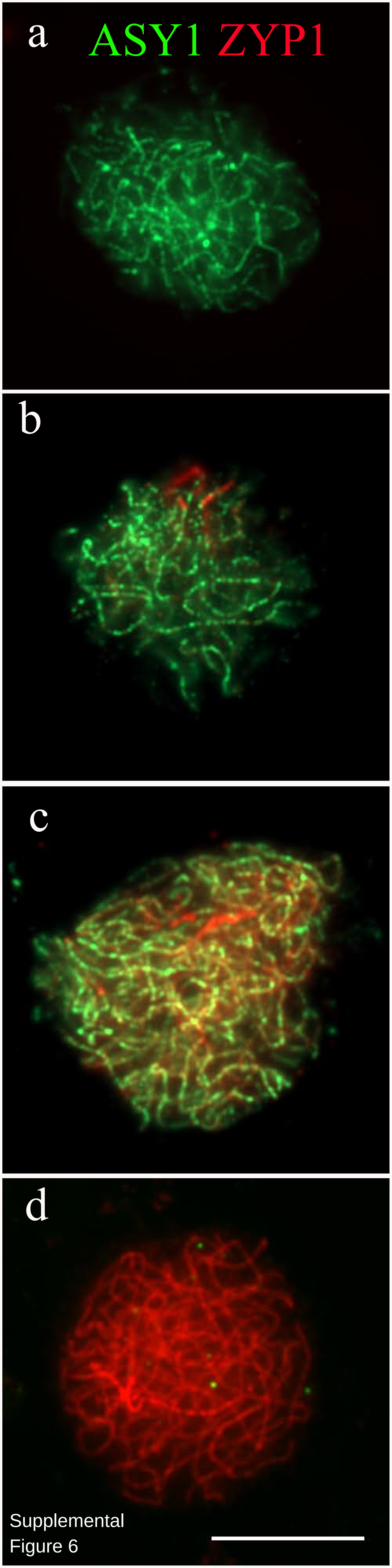


Supplemental
Figure 6

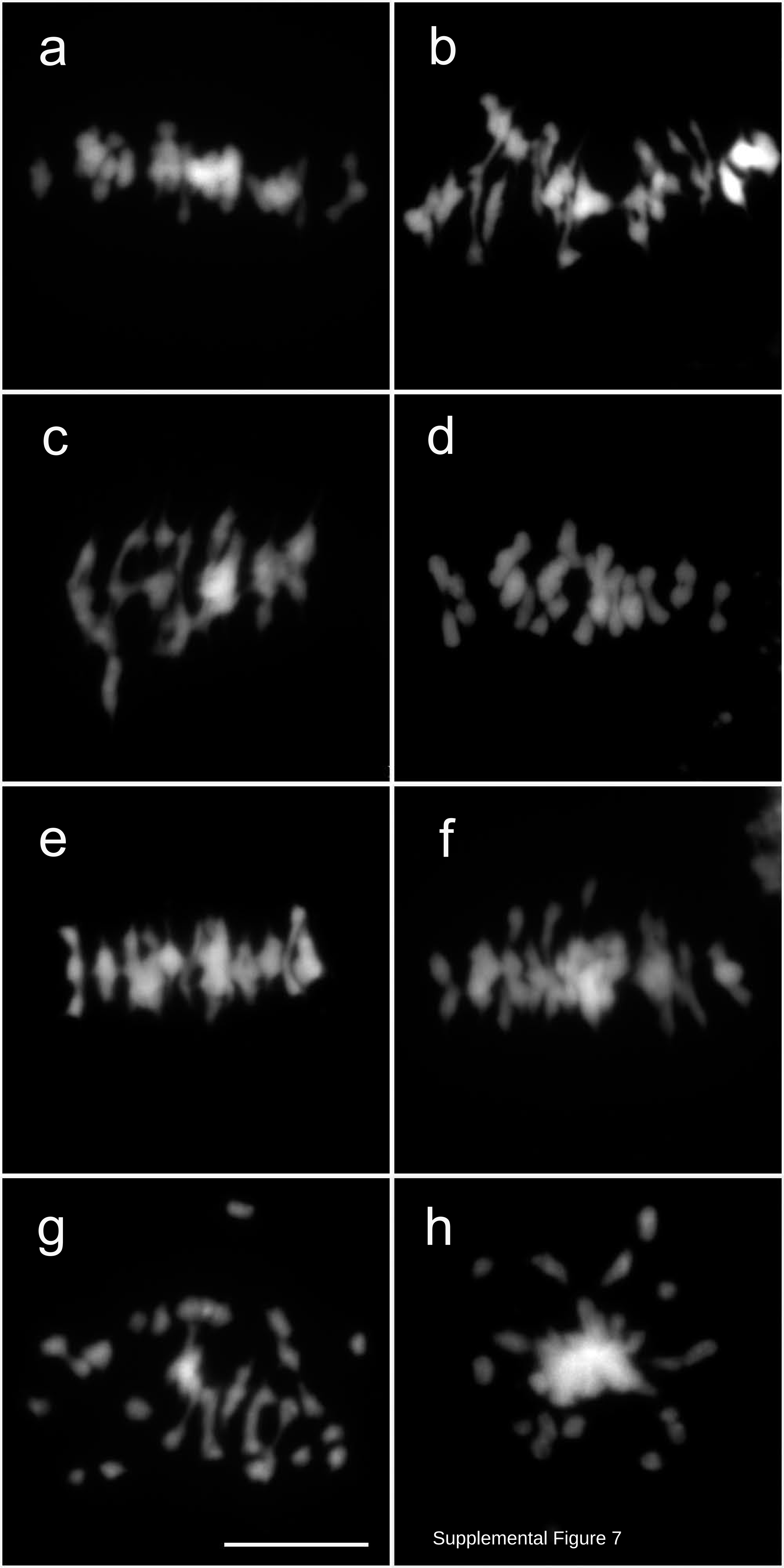


Supplemental Figure 7

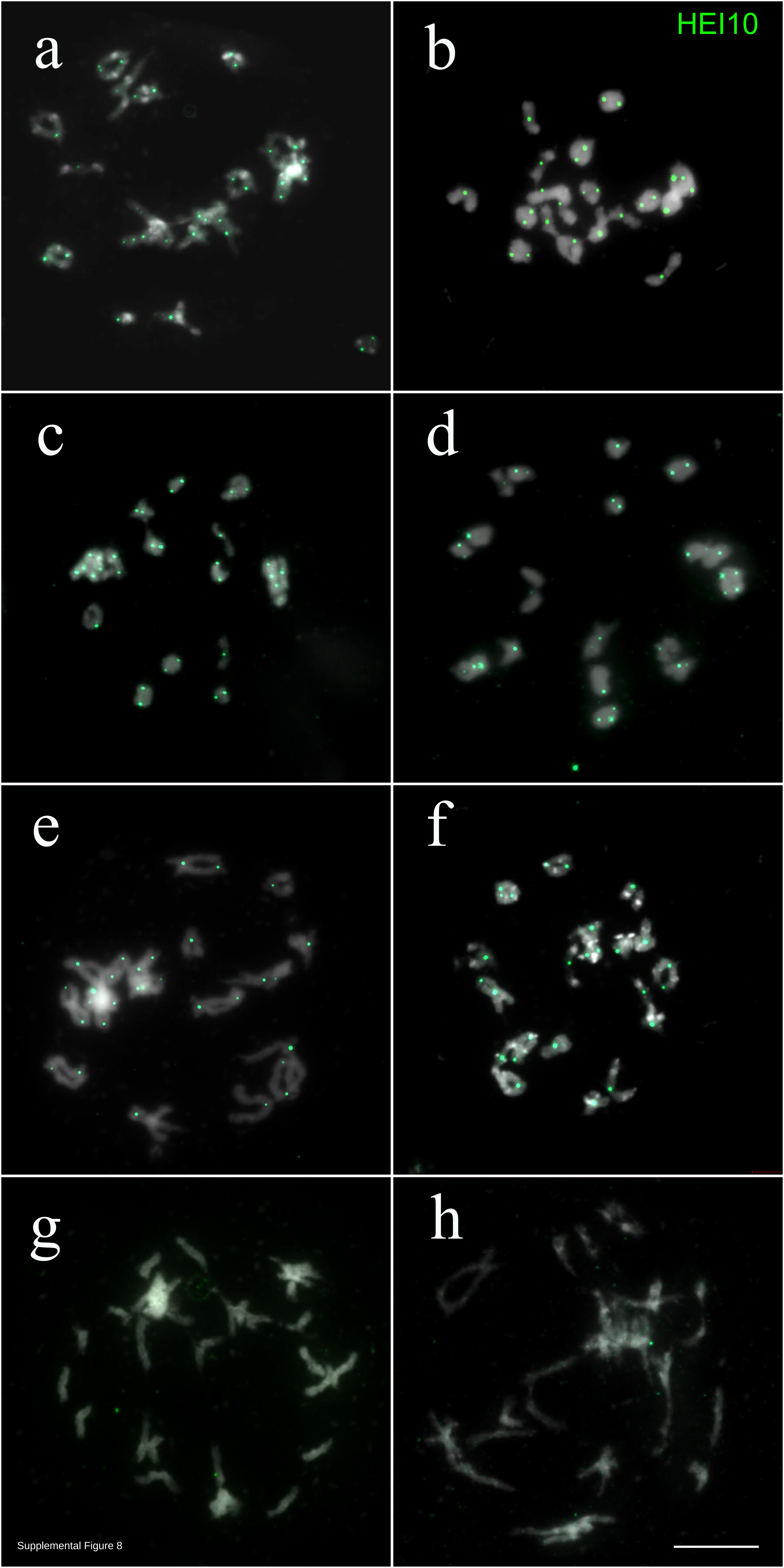

Supplemental Figure 8

**Supplementary Table 1**: ***MSH4* homologs in *Brassica* species**. Summary of the hits obtained from querying *Ath.MSH4* against *Brassica* genomes.

| Organism | Genome | ID | Gene model (version) | Number of exons | Identity with query |
|---|---|---|---|---|---|
| *A. thaliana* | | *Ath.MSH4* | At4g17380 | 24 | 100% |
| *B. napus* | A | *BnaA.MSH4*[1] | BnaA08g08260D (V5) / BnaA08g07470.1D2 (V8.1) | 24 | 91.3% |
| *B. napus* | A | | BnaA08g08250D[1] | 4 | 91.8% |
| *B. napus* | C | *BnaC.MSH4*[1] | BnaCnng35120D (V5) / BnaC02g15090.1D2 (V8.1)[2] | 24 | 89.4% |
| *B. rapa* | A | *Bra.MSH4* | Bra021052 (V1.5) | 24 | 91.2% |
| *B. oleracea* | C | *Bol.MSH4* | Bo8g048310 (V2.1) | 24 | 91.1% |

[1]: Sequence alignments confirmed that *BnaA.MSH4/BnaA08g08260D* and *BnaC.MSH4/BnaCnng35120D* correspond to the two MSH4 genes isolated by Lloyd et al. (2014) by screening a BAC library from *B. napus* cv. *Darmor-bzh*.

[2]: The location of *BnaC.MSH4* is unclear in *B. napus*. *BnaC.MSH4* was first anchored onto a scaffold that was not assigned to a pseudomolecule in the first genome assembly of *B. napus* cv. *Darmor-bzh* (Chalhoub et al., 2014). *BnaC.MSH4* was then assigned and placed on two different chromosomes between the updated assembly of cv. *Darmor-bzh* (BnaC02g15090.1D2 on chromosome C02; Bayer et al., 2017) and the de novo assembly of cv. Tapidor (i.e. BnaC08g39690.1T on chromosome C08; Bayer et al., 2017). These discrepancies indicate either a recurring problem in the assembly of the *BnaC.MSH4* region in *B.napus* cv. *Darmor-bzh* or that the same gene is located on different chromosomes in different varieties (which seems less likely). This notwithstanding, *BnaC.MSH4* is highly related to Bo8g048310 (Figure 1b), which is also located on chromosome C08, and appears as the homoeologous copy of *BnaA.MSH4* (see also Chalhoub et al. 2014).

**Supplementary Table 2: Summed and relative expression of *BnaA.MSH4* and *BnaC.MSH4* in euploid and allohaploid plants combining wild type and mutant alleles for *MSH4***

| Genotypes[1] | # of plants | Quantification of total gene expression[2] | | Relative contribution [3] of | |
|---|---|---|---|---|---|
| | | $2^{-\Delta\Delta Cq}$ | SD | *BnaC.MSH4* | *BnaA.MSH4* |
| *Darmor* | 1 | nd | nd | 83% | 17% |
| *Yudal* | 1 | nd | nd | 72% | 28% |
| *Tanto* | 2 | 0.56 | 0.08 | 78% | 22% |
| $A^+A^+C^+C^+$ | 3 | 1.00 | 0.06 | 79% | 21% |
| $A^1A^1\ C^+C^+$ | 3 | 0.62 | 0.26 | 89% | 11% |
| $A^+A^+C^1C^1$ | 3 | 0.36 | 0.20 | 72% | 28% |
| $A^+A^1C^1C^1$ | 3 | 0.57 | 0.18 | 81% | 19% |
| $A^1A^1C^1C^1$ | 3 | 0.51 | 0.15 | 78% | 22% |
| $A^+C^+$ | 3 | 1.00 | 0.13 | 79% | 21% |
| $A^1C^+$ | 3 | 1.94 | 1.14 | 79% | 21% |
| $A^+C^1$ | 1 | 1.52 | | 75% | 25% |
| $A+C^2$ | 1 | 0.49 | | 76% | 24% |
| $A^2C^1$ | 1 | 1.17 | | nd | nd |

[1]: *Darmor*, *Yudal* and *Tanto* are three wild type cultivars of *B. napus*. The first five genotypes below were identified in the self-fertilized progeny of one F1 hybrid that combined Wild Type *MSH4* alleles (A+ or C+) with mutant *msh4* alleles ($A^1$ or $C^1$; see text for details); these plants are euploids, i.e. they contain the normal number of chromosomes for *B. napus* (2n=38). The five last genotypes are allohaploids (n=19) isolated by microspore culture from two different F1 hybrids that combined Wild Type and mutant alleles for *MSH4*.

[2]: The summed expression of *BnaA.MSH4* and *BnaC.MSH4* was determined by real-time polymerase chain reaction (qPCR). It is expressed as the normalized fold change ($2^{-\Delta\Delta Cq}$) of the target sample relative to a reference sample ($A^+A^+C^+C^+$ for euploids, $A^+C^+$ for allohaploids).

[3]: The relative expression of *BnaA.MSH4* and *BnaC.MSH4* was determined by pyrosequencing. It is presented in the form of percentage.

**Supplementary Table 3. Bivalent and chiasma frequencies between homoeologous chromosomes observed in *Brassica napus* allohaploids.**

| Genotype | Bivalent frequency [1] | Chiasmata frequency [1] | Sample size |
|---|---|---|---|
| $A^{+}C^{+}$ | 5.47 ± 1.4 | 6.4 ± 1.9 | N=90 |
| $A^{1}C^{+}$ | 5 ± 1.7 | 5.4 ± 1.9 | N=112 |
| $A^{+}C^{1}$ | 2.1 ± 1.3 | 2.2 ± 1.5 | N=34 |
| $A^{+}C^{2}$ | 2.3 ± 0.6 | 2.7 ± 0.2 | N=3 |
| $A^{1}C^{1}$ | 1.1 ± 2.7 | 0.7 ± 1.1 | N=34 |
| $A^{1}C^{2}$ | 1.3 ±1.6 | 1.3 ± 1.6 | N=18 |

[1]: Data expressed as mean ± SD

[2]: total number of Pollen Mother Cells observed at Metaphase I (across all plants showing the same genotype).

**Supplementary Table4: Primers used in the study.**

| Name | Sequence 5'-3' | Tm (ºC) |
|---|---|---|
| Q_UBC21F1 | CCTCTGCAGCCTCCTCAAGT | 62 |
| Q_UBC21R1 | GTGTACATGTGTGCCATTGA | 62 |
| Q_MSH4F1 | GATTGTCACCAACCACTCACTCAC | 62 |
| Q_MSH4R1 | CCTCTGCAGCCTCCTCAAGT | 62 |
| PS_MSH41F | TGCTCAGATTGGCTGCTATGT | 62 |
| PS_MSH4S | CAACCACACGCATAGT | 62 |
| PS_MSH4R | TTCTTGTGAATATGCGGTCAAC | 62 |
| T_MSH4AF1 | TTGACCAAAAAATCATCATCGG | 63 |
| T_MSH4AR1 | TGCTATGCTGTAATGAATCCAGAT | 63 |
| T_MSH4CF1 | TTGTCCAAAAAATGTCCTCATCGA | 63 |
| T_MSH4CR1 | CTCAAGACACAAGGATGTATTACGA | 63 |
| Amsh4-57F | TGATGGCAAATGAACTTTCC | 54.5 |
| Amsh4-57R | GCGAGTGGGATGAAGTC | 54.5 |
| Cmsh4-71F | GACGGGTTAGCAATGCCATG | 59.4 |
| Cmsh4-71R | CATTCGGGTAGATGGTTGCC | 59.4 |
| Cmsh4-44F | ATCGCTGAAAGGTACTTGAC | 52 |
| Cmsh4-44R | CAGAACCCATGCGACT | 52 |

**Supplementary Table5: List of cleaved amplified polymorphic markers for identification of *msh4* alleles**

| Allele | Primer pair | Enzyme | restriction fragment size (bp) | |
|---|---|---|---|---|
| | | | **Wt** | **mutant** |
| *BnaA.msh4-1* | Amsh4-57F<br>Amsh4-57R | HpaII | 139+19 | 158 |
| *BnaC.msh4-1* | Cmsh4-71F<br>Cmsh4-71R | NcoI | 196+16 | 212 |
| *BnaC.msh4-2* | Cmsh4-44F<br>Cmsh4-44R | HphI | 177+32 | 209 |

**Supplementary Table 6 : Accession numbers for all proteins**

| Species | common name | Remark | NCBI Reference Sequence | other source |
|---|---|---|---|---|
| **Angiosperms** | | | | |
| *Aegilops tauschii* | | | KQK16204.1 | |
| *Amaranthus hypochondriacus* | | | | https://phytozome.jgi.doe.gov/pz/portal.html#!gene?search=1&crown=1&detail=1&method=0&searchText=transcriptid:32828650 |
| *Amborella trichopoda* | | | ERN00139.1 | |
| *Ananas comosus* | pineapple | | OAY71581.1 | |
| *Aquilegia coerulea* | | | | https://phytozome.jgi.doe.gov/pz/portal.html#!gene?search=1&crown=1&detail=1&method=0&searchText=transcriptid:33070744 |
| *Arabidopsis lyrata* | | | XP_020875333.1 | |
| *Arabidopsis thaliana* | | | NP_193469.2 | |
| *Asparagus officinalis* | garden asparagus | | XP_020263533.1 | |
| *Beta vulgaris* | beet | | XP_010667038.1 | |
| *Brachipodium distachion* | | | XP_003563107.2 | |
| *Brassica napus* | oilseed rape / canola | homoeolog 2 | CDY59530.1 | |
| *Brassica napus* | oilseed rape / canola | homoeolog 1 | XP_013655945.1 | |
| *Brassica oleracea* | kale, cabbage, broccoli, cauliflower | | XP_013605251.1 | |
| *Brassica rapa* | turnip, chinese cabbage, pak choi | | XP_009108319.1 | |
| *Brassica rapa* | | fractionated copy | | |
| *Cajanus cajan* | pigeon pea | | XP_020217246.1 | |

| | | | | |
|---|---|---|---|---|
| *Capsella rubela* | | | XP_006282563.1 | |
| *Cephalotus follicularis* | Australian/Albanian pitcher | | GAV72825.1 | |
| *Chenopodium quinoa* | quinoa | homoeolog 1 | XP_021767980.1 | |
| *Chenopodium quinoa* | | homoeolog 2 | XP_021757807.1 | |
| *Cicer arietinum* | chickpea | | XP_004511707.1 | |
| *Citrullus lanatus* | watermelon | | | http://bioinformatics.psb.ugent.be/plaza/versions/plaza_v3_dicots/genes/view/CL01G05640 |
| *Citrus Clementina* | clementine | | XP_006440279.1 | |
| *Citrus sinensis* | sweet orange | | XP_006477159.1 | |
| *Coffea canephora* | coffee tree | | CDP09986.1 | |
| *Cucumis melo* | Muskmelon | | XP_008437055.1 | |
| *Cucumis sativus* | Cucumber | | XP_011656157.1 | |
| *Cynara cardunculus* | cardoon | | KVH95538.1 | |
| *Daucus Carota* | carot | | XP_017235752.1 | |
| *Dendrobium catenatum* | | | XP_020684536.1 | |
| *Elaeis guineensis* | African oil palm | | XP_010933638.1 | |
| *Eucalyptus grandis* | flooded / rose gum | | XP_010055480.1 | |
| *Fragaria vesca* | wild strawberry | | XP_004301184.1 | |
| *Genlisea aurea* | | | EPS66790.1 | |
| *Glycine max* | soybean | | XP_003538746.1 | |
| *Glycine max* | | | KHN44112.1 | |
| *Gossypium arboreum* | tree cotton | | XP_017634256.1 | |
| *Gossypium hirsutum* | cotton | homoeolog 1 | XP_016733710.1 | |
| *Gossypium hirsutum* | | homoeolog 2 | XP_016665363.1 | |
| *Gossypium raimondii* | | | XP_012439465.1 | |
| *Helianthus annuus* | sunflower | | OTG21795.1 | |
| *Herrania umbratica* | | | XP_021290548.1 | |

| | | | | |
|---|---|---|---|---|
| *Hevea brasiliensis* | rubber tree | | XP_021655095.1 | |
| *Hordeum vulgare* | barley | | AFP73610.1 | |
| *Ipomoea nil* | morning glory | | XP_019173490.1 | |
| *Jatropha curcas* | bubble bush | | XP_012070017.1 | |
| *Juglans regia* | common walnut | | XP_018819253.1 | |
| *Kalanchoe laxiflora* | Milky Widow's Thrill | | | https://phytozome.jgi.doe.gov/pz/portal.html#!gene?search=1&crown=1&detail=1&method=0&searchText=transcriptid:32566639 |
| *Lactuca sativa* | lettuce | | | https://phytozome.jgi.doe.gov/pz/portal.html#!gene?search=1&crown=1&detail=1&er=1&method=0&searchText=transcriptid:38917883 |
| *Linum usitatissimum* | flax | | | https://phytozome.jgi.doe.gov/pz/portal.html#!gene?search=1&crown=1&detail=1&method=0&searchText=transcriptid:23176466 |
| *Linum usitatissimum* | flax | | | https://phytozome.jgi.doe.gov/pz/portal.html#!gene?search=1&crown=1&detail=1&method=0&searchText=transcriptid:23177215 |
| *Lotus japonicus* | | | | http://bioinformatics.psb.ugent.be/plaza/versions/plaza_v3_dicots/genes/view/LJ2G034300 |
| *Macleaya cordata* | plume poppy | | OVA05906.1 | |
| *Malus domestica* | apple tree | | XP_008345359.1 | |
| *Malus domestica* | apple tree | | XP_008345359.1 | |
| *Manihot esculenta* | cassava | | OAY60056.1 | |
| *Medicago truncatula* | | | XP_003611337.2 | |
| *Mimulus guttatus* | monkeyflower | | | https://phytozome.jgi.doe.gov/pz/portal.html#!gene?search=1&crown=1&detail=1&method=0&searchText=transcriptid:28938773 |
| *Momordica charantia* | bitter melon | | XP_022154824.1 | |
| *Musa acuminata* | banana | tandem duplicate | XP_009411922.1 | |
| *Musa acuminata* | | tandem duplicate | XP_009411922.1 | |
| *Nelumbo nucifera* | sacred lotus | | XP_010258855.1 | |
| *Oryza sativa indica* | asian rice | | | http://bioinformatics.psb.ugent.be/plaza/versions/plaza_v3_monocots/genes/view/OSINDICA_07G23690 |

| | | | | |
|---|---|---|---|---|
| *Oryza sativa japonica* | asian rice | | XP_015645440.1 | http://bioinformatics.psb.ugent.be/plaza/versions/plaza_v3_dicots/genes/view/OS07G30240 |
| *Panicum hallii* | Hall's panicgrass | | | https://phytozome.jgi.doe.gov/pz/portal.html#!gene?search=1&crown=1&detail=1&method=0&searchText=transcriptid:32489842 |
| *Phalaenopsis equestris* | | | XP_020585216.1 | |
| *Phoenix dactylifera* | date palm | | XP_008796122.1 | |
| *Populus trichocarpa* | poplar | | PNT54745.1 | http://bioinformatics.psb.ugent.be/plaza/versions/plaza_v3_dicots/genes/view/PT01G15620 |
| *Populus trichocarpa* | poplar | Fractionated copy | | http://bioinformatics.psb.ugent.be/plaza/versions/plaza_v3_dicots/genes/view/PT03G07860 |
| *Prunus persica* | peach tree | | XP_007210902.1 | |
| *Pyrus x bretschneideri* | pear | | XP_009334525.1 | |
| *Raphanss sativus* | radish | | XP_018447151.1 | |
| *Ricinus communis* | castor bean | | XP_015581765.1 | |
| *Salix purpurea* | purple willow | | | https://phytozome.jgi.doe.gov/pz/portal.html#!gene?search=1&crown=1&detail=1&method=0&searchText=transcriptid:31422618 |
| *Sesamum indicum* | sesame | | XP_011081405.1 | |
| *setaria italica* | Foxtail millet | | KQL25810.1 | |
| *Setaria viridis* | green foxtail | | | https://phytozome.jgi.doe.gov/pz/portal.html#!gene?search=1&crown=1&detail=1&method=0&searchText=transcriptid:32637916 |
| *Solanum lycopersicum* | tomato | | XP_004244563.1 | |
| *Solanum tuberosum* | potato | | XP_006358345.1 | |
| *Sorghum bicolor* | sorghum | | | https://phytozome.jgi.doe.gov/pz/portal.html#!gene?search=1&crown=1&detail=1&method=0&searchText=transcriptid:37953771 |
| *Spinacia oleracea* | spinach | | XP_021855422.1 | |
| *Spirodela polyrhiza* | Common duckmeat | | | https://phytozome.jgi.doe.gov/pz/portal.html#!gene?search=1&crown=1&detail=1&method=0&searchText=transcriptid:31522044 |
| *Theobroma cacao* | cacao | | EOY24295.1 | |
| *Zea mays* | maize | | APZ84449.1 | |
| *Ziziphus jujuba* | Common Jujube | | XP_015896598.1 | |
| *Zostera marina* | eelgrass | | KMZ57700.1 | |

| Fungi | | | | |
|---|---|---|---|---|
| *Agaricus bisporus* | | | | https://genome.jgi.doe.gov/cgi-bin/dispGeneModel?db=Agabi_varbisH97_2&id=187616 |
| *Ajellomyces dermatitidis* | | | | https://www.uniprot.org/uniprot/F2TIP8 |
| *Allomyces macrogynus* | | | KNE71791 | http://fungi.ensembl.org/Allomyces_macrogynus_atcc_38327/Transcript/ProteinSummary?db=core;g=AMAG_16098;r=supercont3.47:254149-257748;t=KNE71791;tl=5sNtdTv0Vl7GEZe0-18523558-450668881 |
| *Alternaria alternata* | | | XP_018386533.1 | |
| *Amanita thiersii* | | | | https://genome.jgi.doe.gov/cgi-bin/dispGeneModel?db=Amath1&id=150394 |
| *Arthrobotrys oligospora* | | | | https://www.uniprot.org/uniprot/G1X7G6 |
| *Ascoidea rubescens* | | | XP_020048208.1 | |
| *Aspergillus glaucus* | | | XP_022403503.1 | |
| *Aureobasidium subglaciale* | | | XP_013346404.1 | |
| *Batrachochytrium dendrobatidis* | | | OAJ39063.1 | |
| *Baudoinia panamericana* | | | XP_007681481.1 | |
| *Bipolaris oryzae* | | | XP_007691950.1 | |
| *Blastocladiella britannica* | | | | https://genome.jgi.doe.gov/cgi-bin/dispGeneModel?db=Blabri1&id=137578 |
| *Botrytis cinerea* | | | XP_024546274.1 | |
| *Candida albicans* | | | XP_718245.2 | |
| *Candida orthopsilosis* | | | XP_003865918.1 | |
| *Candida_arabinofermentans* | | | ODV88025.1 | |
| *Catenaria anguillulae* | | | ORZ32677.1 | |
| *Cercospora beticola* | | | XP_023450075.1 | |
| *Cladonia grayi* | | | | https://genome.jgi.doe.gov/cgi-bin/dispGeneModel?db=Clagr3&id=9429 |

| | | | | |
|---|---|---|---|---|
| *Colletotrichum orchidophilum* | | | XP_022475615.1 | |
| *Coprinopsis cinerea* | | | | https://genome.jgi.doe.gov/cgi-bin/dispGeneModel?db=Copci1&id=11167 |
| *Cryptococcus neoformans* | | | | https://genome.jgi.doe.gov/cgi-bin/dispGeneModel?db=Cryne_H99_1&id=4689 |
| *Cyphellophora europaea* | | | XP_008719429.1 | |
| *Dactylellina haptotyla* | | | | https://www.uniprot.org/uniprot/S8BYD4 |
| *Diplocarpon rosae* | | | PBP25854.1 | |
| *Diplocarpon rosae* | | | PBP17310.1 | |
| *Eremothecium cymbalariae* | | | XP_003647962.1 | |
| *Fusarium graminearum* | | | XP_011324276.1 | |
| *Glarea lozoyensis* | | | XP_008083213.1 | |
| *Hanseniaspora osmophila* | | | OEJ85393.1 | |
| *Hortaea werneckii* | | | OTA24137.1 | |
| *Hortaea werneckii* | | | OTA35336.1 | |
| *Hyphopichia burtonii* | | | XP_020076843.1 | |
| *Kazachstania africana* | | | XP_003956462.1 | |
| *Kazachstania naganishii* | | | XP_022465397.1 | |
| *Kluyveromyces dobzhanskii* | | | CDO95828.1 | |
| *Kluyveromyces lactis* | | | XP_455582.1 | |
| *Kluyveromyces marxianus* | | | XP_022674546.1 | |
| *Lichtheimia corymbifera* | | | | https://genome.jgi.doe.gov/cgi-bin/dispGeneModel?db=Liccor1&id=6941 |
| *Lobosporangium transversale* | | | XP_021882207.1 | |

| | | | | |
|---|---|---|---|---|
| *Lodderomyces elongisporus* | | | XP_001527607.1 | |
| *Marssonina brunnea* | | | XP_007292754.1 | |
| *Melampsora larici-populina* | | | | https://genome.jgi.doe.gov/cgi-bin/dispGeneModel?db=Mellp2_3&id=106496 |
| *Meliniomyces bicolor* | | | XP_007780362.1 | |
| *Moesziomyces aphidis* | | | | https://genome.jgi.doe.gov/cgi-bin/dispGeneModel?db=Moeaph1&id=1263 |
| *Mortierella elongata* | | | KFH69849.1 | |
| *Mucor circinelloides* | | | | https://genome.jgi.doe.gov/cgi-bin/dispGeneModel?db=Mucci2&id=42353 |
| *Naumovozyma castellii* | | | XP_003675758.1 | |
| *Naumovozyma dairenensis* | | | XP_003671356.2 | |
| *Neolecta irregularis* | | | | https://www.uniprot.org/uniprot/A0A1U7LMA1 |
| *Neurospora crassa* | | | XP_001728471.2 | |
| *Neurospora tetrasperma* | | | EGZ76821.1 | |
| *Oidiodendron maius* | | | KIN03945.1 | |
| *Pestalotiopsis fici* | | | XP_007827801.1 | |
| *Phanerochaete chrysosporium* | | | | https://genome.jgi.doe.gov/cgi-bin/dispGeneModel?db=Phchr2&id=3013195 |
| *Phialocephala scopiformis* | | | XP_018076977.1 | |
| *Phycomyces blakesleeanus* | | | | https://genome.jgi.doe.gov/cgi-bin/dispGeneModel?db=Phybl2&id=64866 |
| *Protomyces lactucaedebilis* | | | | https://genome.jgi.doe.gov/cgi-bin/dispGeneModel?db=Prola1&id=409956 |
| *Pseudocercospora fijiensis* | | | XP_007929281.1 | |
| *Pseudogymnoascus verrucosus* | | | XP_018132570.1 | |

| | | | | |
|---|---|---|---|---|
| *Pseudozyma antarctica* | | | | https://genome.jgi.doe.gov/cgi-bin/dispGeneModel?db=Psean1_1&id=78532 |
| *Puccinia graminis* | | | | https://genome.jgi.doe.gov/cgi-bin/dispGeneModel?db=Pucgr2&id=12408 |
| *Pycnoporus cinnabarinus* | | | | https://genome.jgi.doe.gov/cgi-bin/dispGeneModel?db=Pycci1&id=5650 |
| *Pyronema omphalodes* | | | | https://www.uniprot.org/uniprot/U4KXX1 |
| *Ramularia collo-cygni* | | | XP_023625185.1 | |
| *Rhizopus delemar* | | | | https://genome.jgi.doe.gov/cgi-bin/dispGeneModel?db=Rhior3&id=10733 |
| *Rhizopus delemar* | | | | https://genome.jgi.doe.gov/cgi-bin/dispGeneModel?db=Rhior3&id=10734 |
| *Saccharomyces bayanus* | | | AACA01000011.1 | |
| *Saccharomyces boulardii* | | | | https://fungi.ensembl.org/Saccharomyces_sp_boulardii__gca_001413975/Transcript/ProteinSummary?db=core;g=AB282_01824;r=VI:125552-128188;t=KQC44160;tl=jFDFbWdP0QnRiFat-18707692-535631528 |
| *Saccharomyces cerevisiae* | | | NP_116652.1 | |
| *Saccharomyces enbayanus* | | | | https://fungi.ensembl.org/Saccharomyces_eubayanus_gca_001298625/Transcript/ProteinSummary?db=core;g=DI49_1650;r=6:115436-118072;t=KOG99815;tl=7aRuGgLrG4P82TP0-18707675-535619505 |
| *Saccharomyces eubayanus* | | | XP_018222533.1 | |
| *Saccharomyces kudriavzevii* | | | | |
| *Saccharomyces mikatae* | | | AABZ01000312.1 | |
| *Saitoella complicata* | | | | https://www.uniprot.org/uniprot/A0A0E9NCE7 |
| *Sclerotinia borealis* | | | ESZ98411.1 | |
| *Sclerotinia sclerotiorum* | | | APA06952.1 | |
| *Sordaria macrospora* | | | XP_003352637.1 | |
| *Sphaerulina musiva* | | | XP_016758040.1 | |

| | | | | |
|---|---|---|---|---|
| *Sporisorium reilianum* | | | | https://genome.jgi.doe.gov/cgi-bin/dispGeneModel?db=Spore1&id=4182 |
| *Tetrapisispora blattae* | | | XP_004181806.1 | |
| *Tetrapisispora phaffii* | | | XP_003685140.1 | |
| *Tilletiaria anomala* | | | | https://genome.jgi.doe.gov/cgi-bin/dispGeneModel?db=Tilan2&id=257515 |
| *Torulaspora delbrueckii* | | | XP_003680201.1 | |
| *Trichosporon chiarellii* | | | | https://genome.jgi.doe.gov/cgi-bin/dispGeneModel?db=Trich1&id=23288 |
| *Tuber aestivum* | | | | https://www.uniprot.org/uniprot/A0A292Q6H3 |
| *Tuber melanosporum* | | | | https://www.uniprot.org/uniprot/D5GKH6 |
| *Urocystis primulicola* | | | | https://genome.jgi.doe.gov/cgi-bin/dispGeneModel?db=Uropr1&id=219307 |
| *Ustilago maydis* | | | | https://genome.jgi.doe.gov/cgi-bin/dispGeneModel?db=Ustma2_2&id=6671 |
| *Vanderwaltozyma polyspora* | | | XP_001645356.1 | |
| *Verruconis gallopava* | | | XP_016211195.1 | |
| *Wickerhamomyces anomalus* | | | XP_019041758.1 | |
| *Wickerhamomyces ciferrii* | | | XP_011271514.1 | |
| *Xanthoria parietina* | | | | https://genome.jgi.doe.gov/cgi-bin/dispGeneModel?db=Xanpa2&id=1615593 |
| *Zygosaccharomyces rouxii* | | | GAV54848.1 | |
| *Zygosaccharomyces rouxii* | | | GAV49934.1 | |
| *Zymoseptoria tritici* | | | XP_003850619.1 | |
| **Animals** | | | | |
| *Alligator mississipiensis* | Alligator | | XP_019344642.1 | |

| | | | | |
|---|---|---|---|---|
| *Anas platyrhynchos* | Duck | | | https://www.ensembl.org/Anas_platyrhynchos/Transcript/ProteinSummary?db=core;t=ENSAPLP00000014123;tl=umMuh1C4mtNprauo-4621137-705896503 |
| *Anolis carolinensis* | Anole Lizard | | | https://www.ensembl.org/Anolis_carolinensis/Transcript/ProteinSummary?db=core;g=ENSACAG00000010892;r=GL343194.1:3378016-3405609;t=ENSACAT00000011040;tl=umMuh1C4mtNprauo-4621134-705896474 |
| *Apis mellifera* | European honey bee | | XP_026299499.1 | |
| *Astyanax mexicanus* | Cave fish | | XP_015462163.2 | |
| *Bombyx mori* | Domestic silkmoth | | XP_021204461.1 | |
| *Bos taurus* | Cow | | | https://www.ensembl.org/Bos_taurus/Transcript/ProteinSummary?db=core;t=ENSBTAP00000024552;tl=hQuE7z7QGDzB1DIk-4621178-705897779 |
| *Branchiostoma floridae* | Amphioxus | | XP_019619725.1 | fgenesh2_pg.scaffold_897000004 |
| *Canis familiaris* | dog | | | https://www.ensembl.org/Canis_familiaris/Transcript/ProteinSummary?db=core;t=ENSCAFP00000033551;tl=hQuE7z7QGDzB1DIk-4621173-705897721 |
| *Ciona savignyi* | Transparent squirt | | ENSCSAVP00000004950 | |
| *Danio rerio* | Zebra Fish | | XP_009291531.1 | |
| *Daphnia pulex* | Waterflea | | EFX81651.1 | |
| *Ficedula albicollis* | Flycatcher | | | https://www.ensembl.org/Ficedula_albicollis/Transcript/ProteinSummary?db=core;t=ENSFALP00000001414;tl=umMuh1C4mtNprauo-4621133-705896468 |
| *Gadus morhua* | Athlantic cod | | | https://www.ensembl.org/Gadus_morhua/Transcript/Summary?db=core;g=ENSGMOG00000016471;r=GeneScaffold_1262:841386-841874;t=ENSGMOT00000018126;tl=Fxvrk8S5Ndpt787t-4621421-705903065 |

| | | | | |
|---|---|---|---|---|
| *Gallus gallus* | Chicken | | | https://www.ensembl.org/Gallus_gallus/Transcript/ProteinSummary?db=core;t=ENSGALP00000018559;tl=umMuh1C4mtNprauo-4621131-705896444 |
| *Gasterosteus aculeatus* | Stickleback | | | https://www.ensembl.org/Gasterosteus_aculeatus/Gene/Summary?db=core;g=ENSGACG00000007948;r=groupXV:5124334-5125063;t=ENSGACT00000010560;tl=Fxvrk8S5Ndpt787t-4621422-705903043 |
| *Homo sapiens* | Human | | | https://www.ensembl.org/Homo_sapiens/Transcript/ProteinSummary?db=core;t=ENSP00000263187;tl=hQuE7z7QGDzB1DIk-4621175-705897818 |
| *Latimeria chalumnae* | Coelacanth | | XP_014349280.1 | |
| *Lepisosteus oculatus* | Spotted gar | | XP_015205696.1 | |
| *Limulus polyphemus* | | | XP_013777928.1 | |
| *Mus musculus* | Mouse | | | https://www.ensembl.org/Mus_musculus/Transcript/ProteinSummary?db=core;t=ENSMUSP00000005630;tl=hQuE7z7QGDzB1DIk-4621174-705897767 |
| *Myotis lucifugus* | bat | | | https://www.ensembl.org/Myotis_lucifugus/Transcript/ProteinSummary?db=core;t=ENSMLUP00000008419;tl=hQuE7z7QGDzB1DIk-4621180-705897731 |
| *Oncorhynchus mykiss* | Rainbow trout | | XP_021440560.1 | |
| *Oncorhynchus mykiss* | Rainbow trout | fractionated copy | | https://salmobase.org/cgi-bin/gb2/gbrowse_details/rainbow_trout_ncbi?ref=NC_035095.1;start=38408622;end=38432645;name=LOC110498353;class=Sequence;feature_id=772966;db_id=rainbow_trout_annotation%3Adatabase |
| *Oreochromis niloticus* | Tilapia | | XP_005477028.1 | |
| *Ornithorhynchus anatinus* | Platypus | | | https://www.ensembl.org/Ornithorhynchus_anatinus/Transcript/ProteinSummary?db=core;t=ENSOANP00000025217;tl=ictrlDIwrnrvAdvZ-4621466-705903586 |
| *Oryctolagus cuniculus* | Rabit | | | https://www.ensembl.org/Oryctolagus_cuniculus/Transcript/ProteinSummary?db=core;t=ENSOCUP00000013962;tl=hQuE7z7QGDzB1DIk-4621181-705897805 |
| *Oryzias latipes* | Medaka | | XP_020569477.1 | |

| | | | | |
|---|---|---|---|---|
| *Pan troglodytes* | Chimpanzee | | | https://www.ensembl.org/Pan_troglodytes/Transcript/ProteinSummary?db=core;t=ENSPTRP00000084312;tl=umMuh1C4mtNprauo-4621135-705896490 |
| *Parasteatoda tepidariorum* | Common House Spider | | | |
| *Pelodiscus sinensis* | Chinesse shortshell turtle | | XP_025044129.1 | |
| *Poecilia formosa* | Amazon molly | | XP_008398407.1 | |
| *Salmo salar* | Atlantic salmon | | XP_014066431 | |
| *Salmo salar* | Atlantic salmon | fractionated copy | | https://salmobase.org/cgi-bin/gb2/gbrowse_details/salmon_GBrowse_Chr_NCBI?ref=ssa01;start=25683799;end=25694444;name=LOC106586026;class=Sequence;feature_id=2907398;db_id=salmon_annotation%3Adatabase |
| *Salvelinus alpinus* | Arctic char | | XP_023842423.1 | |
| *Strongylocentrotus purpuratus* | Purple sea urchin | | XP_011662355.1 | |
| *Takifugu rubripes* | Fugu | | | https://www.ensembl.org/Takifugu_rubripes/Gene/Summary?db=core;g=ENSTRUG00000018083;r=2:7183999-7195134;t=ENSTRUT00000046458;tl=Fxvrk8S5Ndpt787t-4621423-705902999 |
| *Tetraodon nigroviridis* | Tetraodon | | | https://www.ensembl.org/Tetraodon_nigroviridis/Transcript/Summary?db=core;g=ENSTNIG00000017376;r=10:5576872-5577274;t=ENSTNIT00000020746;tl=Fxvrk8S5Ndpt787t-4621420-705903023 |
| *Xenopus laevis* | African clawed frog | | XP_018112574 | |
| *Xenopus tropicalis* | Western clawed frog | | XP_017949021.1 | |
| *Xiphophorus maculatus* | Platyfish | | XP_014331698.1 | |
| **outgroups** | | | | |
| *Caenorhabditis elegans* | | | NP_495451.1 | |

| | | | | |
|---|---|---|---|---|
| *Chlamydomonas reinhardtii* | | | | http://bioinformatics.psb.ugent.be/plaza/versions/plaza_v3_dicots/genes/view/CR16G00840 |
| *Condrus crispus* | | | | http://plants.ensembl.org/Chondrus_crispus/Transcript/Sequence_Protein?db=core;g=CHC_T00008573001;r=HG002195:34740-37806;t=CDF40516 |
| *Conidiobolus thromboides* | | | | https://genome.jgi.doe.gov/cgi-bin/dispGeneModel?db=Conth1&id=350361 |
| *Cyanidioschyzon merolae* | | | | http://plants.ensembl.org/Cyanidioschyzon_merolae/Transcript/Sequence_Protein?db=core;g=CMK199C;r=11:547550-550294;t=CMK199CT |
| *Galdieria sulphuraria* | | | | http://plants.ensembl.org/Galdieria_sulphuraria/Transcript/Sequence_Protein?db=core;g=Gasu_57130;r=scaf_60:70601-75041;t=EME26709 |
| *Selaginella moellendorffi* | | | | http://plants.ensembl.org/Selaginella_moellendorffii/Transcript/Sequence_Protein?db=core;g=SELMODRAFT_115028;r=GL377615:864082-867676;t=EFJ17259 |
| *Rozella allomycis* | | | EPZ31156.1 | |
| *Selaginella moellendorffi* | | | | http://plants.ensembl.org/Selaginella_moellendorffii/Transcript/Sequence_Protein?db=core;g=SELMODRAFT_88599;r=GL377574:489231-492826;t=EFJ30915 |
| *Physcomitrella patens* | | | XP_001777754.1 | |
| *Sphagnum fallax* | | | | https://phytozome.jgi.doe.gov/pz/portal.html#!gene?search=1&crown=1&detail=1&method=0&searchText=transcriptid:32620337 |
